# Copper chelation reprograms the tumour microenvironment to enhance immune checkpoint blockade therapy in preclinical mesothelioma

**DOI:** 10.64898/2026.07.28.740857

**Authors:** Kofi L.P. Stevens, Naomi Damstra, Claudia Peh, Nicola Principe, Amber-Lee Phung, Joshua Ravensdale, Sophie Wu, Vu Pham, Giulia Castrogiovanni, Louis Boon, Keea R. Inder-Smith, Wen Shuz Yeow, Daryl L. Howard, Ramesh Ram, Abha Chopra, Willem Joost Lesterhuis, Rachael M. Zemek, Benjamin Kong, Andrew Crowe, Delia Nelson, Ross M. Graham, Wee Loong Chin, Orazio Vittorio, Jonathan Chee

## Abstract

Immune checkpoint blockade provides durable benefit to only a minority of patients with pleural mesothelioma. Here we investigated whether targeting copper homeostasis could enhance anti-tumour immunity and improve responses to immune checkpoint blockade. Copper homeostasis genes were upregulated in human mesothelioma, localised to malignant and stromal cells but downregulated in lymphocytes. In murine mesothelioma, these genes were dynamically regulated during anti-CTLA-4 and anti-PD-1 therapy. Treatment with copper chelator triethylenetetramine reduced intratumoural copper, increased cytotoxic lymphocytes, and enriched inflammatory genes in tumour-infiltrating myeloid and lymphoid compartments. Furthermore, copper chelation enhanced anti-CTLA-4 and anti-PD-1 therapy in immunocompetent mesothelioma mouse models, leading to durable tumour control. Our findings identify copper homeostasis as a therapeutic vulnerability in mesothelioma and support clinical evaluation of copper depletion with first-line immune checkpoint blockade.

## INTRODUCTION

Pleural mesothelioma is an aggressive asbestos-related cancer with a five-year survival rate of approximately 10%. Despite the introduction of first-line immune checkpoint blockade (ICB) that targets Programmed cell Death Protein 1 (PD-1) and Cytotoxic T-Lymphocytes Associated protein 4 (CTLA-4), only a minority of patients derive clinical benefit, and responses are rarely durable (*1, 2*). Improving the frequency and duration of responses to ICB is critical for these patients. Tumours with high CD8⁺ T cell infiltration and proinflammatory gene signatures are more likely to respond to ICB (*3, 4*), implying that therapeutic strategies that improve T cell infiltration and function, could augment ICB efficacy.

Copper accumulates in malignant cells at levels exceeding those in normal tissues likely because it is an important co-factor for enzymes that regulate cell proliferation, reactive oxidation species clearance and angiogenesis (*5*). Consistent with this, human and mouse mesothelioma tumours progressively sequester copper (*6, 7*). Beyond supporting tumour growth, we have identified copper as an immunoregulatory signal within the tumour microenvironment (TME). Excess intratumoural copper increases expression of the immune checkpoint molecule programmed death-ligand 1 (PD-L1) via STAT3 signalling, providing a targetable, copper-mediated pathway of tumour immune evasion (*8, 9*). These findings are consistent with the broader concept that malignant cells develop a heightened dependency on copper to sustain proliferation and stress adaptation, potentially limiting copper availability and metabolic fitness within tumour infiltrating immune cells. Therapeutic copper depletion with metal chelators remodelled the myeloid and lymphoid components of the TME, and significantly improved responses to a different class of immunotherapy (anti-GD2) in another cancer model (*8*). Together, these observations prompted us to investigate whether targeting copper metabolism with triethylenetetramine (TETA, marketed as Cuprior) can remodel the mesothelioma TME, alter immune suppression and improve responses to anti-CTLA-4 and anti-PD-1 ICB.

In this study, we report enrichment of copper homeostasis genes in tumour cells relative to immune cells within mouse and human mesothelioma datasets. TETA reduces intratumoural copper, alters both lymphoid and myeloid compartments and increases cytotoxic CD8^+^ T cell and NK cells. We demonstrate in two immunocompetent mesothelioma murine models that copper chelation with TETA is an effective strategy to improve anti-CTLA-4 and anti-PD-1 therapy. Together, these findings provide crucial evidence to support the clinical testing of combination therapy for patients with mesothelioma.

## RESULTS

### Copper homeostasis gene programs are upregulated in human mesothelioma and localise to malignant cells and fibroblasts

As tumours have elevated copper levels in comparison to normal tissue (*10*), we hypothesised that mesothelioma tumours would be enriched in copper homeostasis genes compared to non-cancerous tissue. We interrogated publicly available human mesothelioma gene expression datasets from the TCGA and GSE51024 (*11*) for expression of genes that regulate copper import, export, intracellular buffering and cuproenzyme activity, henceforth referred to as copper homeostasis genes (Fig. 1A) (*12*). 76 out of 121 copper homeostasis genes were significantly enriched in mesothelioma samples compared to non-cancerous tissues (table S1). This included *SLC31A1*, which encodes for the major high affinity copper importer, and *ATP7A*, which encodes a copper efflux pump (Fig. 1B). In the public TCGA-MESO cohort, the copper homeostasis gene set was not associated with overall survival at the aggregate level, but individual genes showed divergent associations in univariate analysis (table S2). For example, high expression of genes such as *ATP7A* and lower expression of others such as *SLC31A1* were each associated with improved survival (Fig. 1C). We next interrogated publicly available human mesothelioma single cell data (*13*) (Giotti et al., n=12; MOSAIC-MESO n=10) to visualise expression of copper homeostasis genes within the TME (Fig. 1D, fig. S1A, B). Single-cell expression showed that copper homeostasis genes were preferentially expressed in tumour cells and fibroblasts with comparatively low expression in immune cells such as T cells, B cells and NK cells (Fig. 1E, fig. S1C). Consistently, GSEA on ranked differential expression statistics demonstrated significant downregulation of the copper gene homeostasis set in T cells, B cells, NK cells (Fig. 1F, fig. S1D). Pseudobulk analysis showed that key copper transport and metabolism genes such as *ATP7B*, *MT3* and *STEAP4* were significantly upregulated in tumour cells, whereas *MOXD1*, *LOXL1* and *LOXL2* were enriched in fibroblasts (fig. S1E). Overall, copper homeostasis gene upregulation in these human mesothelioma samples is largely tumour-and fibroblast-derived, with minimal immune cell contribution, suggesting elevated copper demand in malignant cells.

**Fig. 1.**
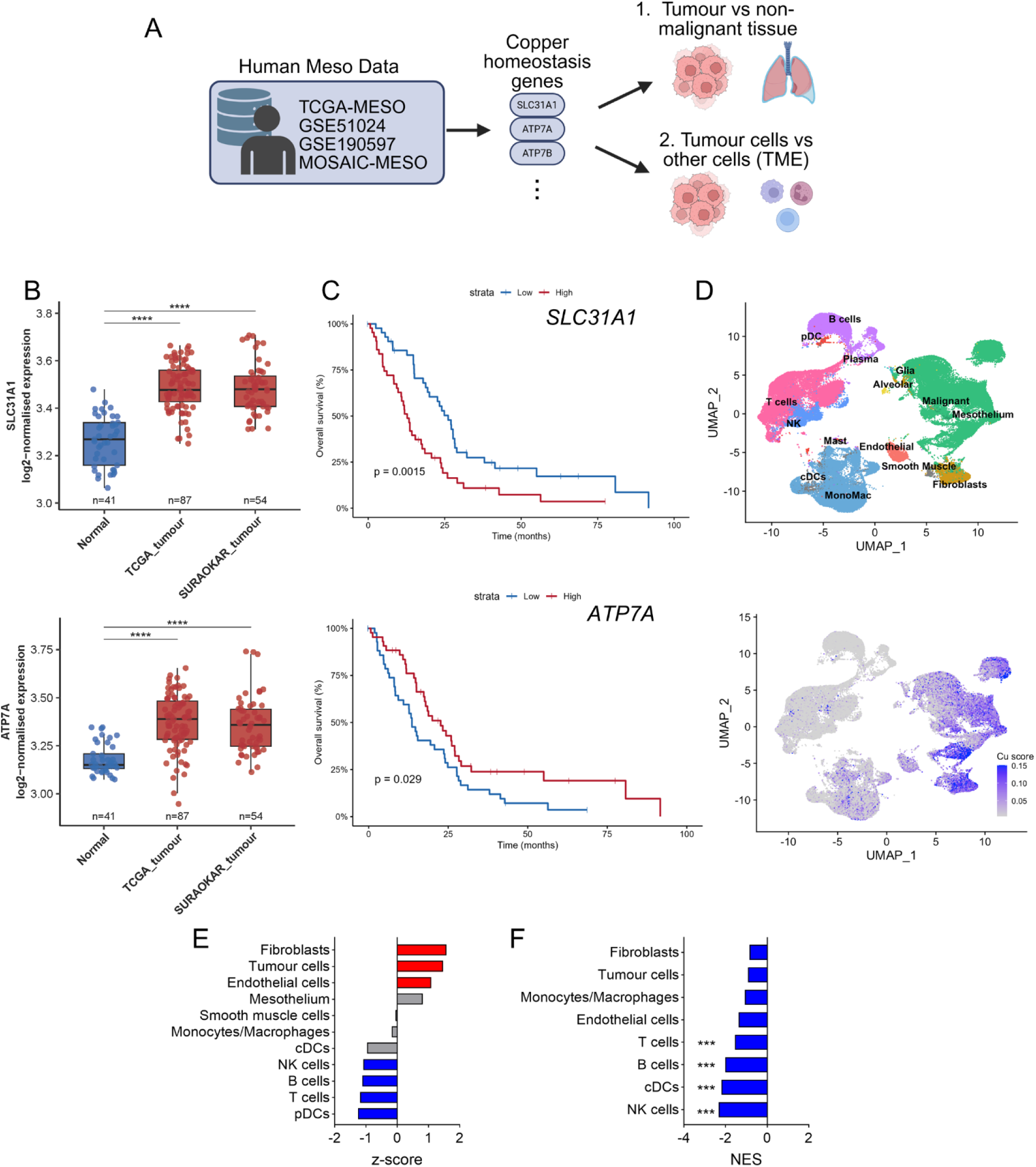
Copper homeostasis gene programs are upregulated in human mesothelioma and localise to malignant and stromal cells. (**A**) Schematic depicting analysis of publicly available human mesothelioma datasets, created with BioRender.com. (**B**) Log2-normalised expression of Cu transport genes from two mesothelioma cohorts compared to normal tissue. Differences in gene expression between cohorts were assessed using the Wilcoxon rank-sum test, with Benjamini–Hochberg correction for multiple comparisons. (**C**) Kaplan-Meier survival curves of copper transport genes stratified by high or low expression using the cohort median cut-offs. Survival differences assessed by Log-rank test. (**D**) UMAP projection of single-cell RNA-seq data from human mesothelioma samples (n = 12), annotated with major cell populations (top) and overlaid with a per-cell copper homeostasis gene-set score (bottom). (**E**) Average expression of copper homeostasis gene set for indicated cell subsets, shown as z-scores. (**F**) Normalised enrichment scores (NES) from GSEA of copper homeostasis gene sets across cell subsets. Cell subsets significantly enriched for copper homeostasis genes are indicated by asterisks. *** padj < 0.005, **** padj < 0.001.

### Murine mesothelioma recapitulates copper homeostasis gene expression in tumour and fibroblasts

We previously utilised murine mesothelioma models to assess the anti-cancer effects of copper chelators (*6*). Mesothelioma cell lines upregulated PD-L1 with the addition of copper in vitro (fig S2A, B), consistent with prior studies in other mouse tumour cell lines (*9*). We analysed previously published single cell RNAseq data from ex vivo murine AB1 mesothelioma tumours (Fig. 2A) (*14*), and observed a similar pattern of copper homeostasis genes preferentially expressed in tumour cells and fibroblasts rather than immune cell populations (Fig. 2B, 2C). Pseudo-bulk analysis showed that genes such as *Loxl3*, *Mt2* and *Steap1* were significantly upregulated in tumour cells, whereas *Lox, Loxl1, Loxl2, Mt1, Mt2, Sod3* and *Steap2* were enriched in fibroblasts (Fig. 2D). Consistent with this, there was significant downregulation of the copper genes in T cells and NK cells (Fig. 2E), indicating that the murine model recapitulates the pattern seen in human mesothelioma where copper homeostasis gene expression is concentrated in malignant rather than immune cells.

**Fig. 2.**
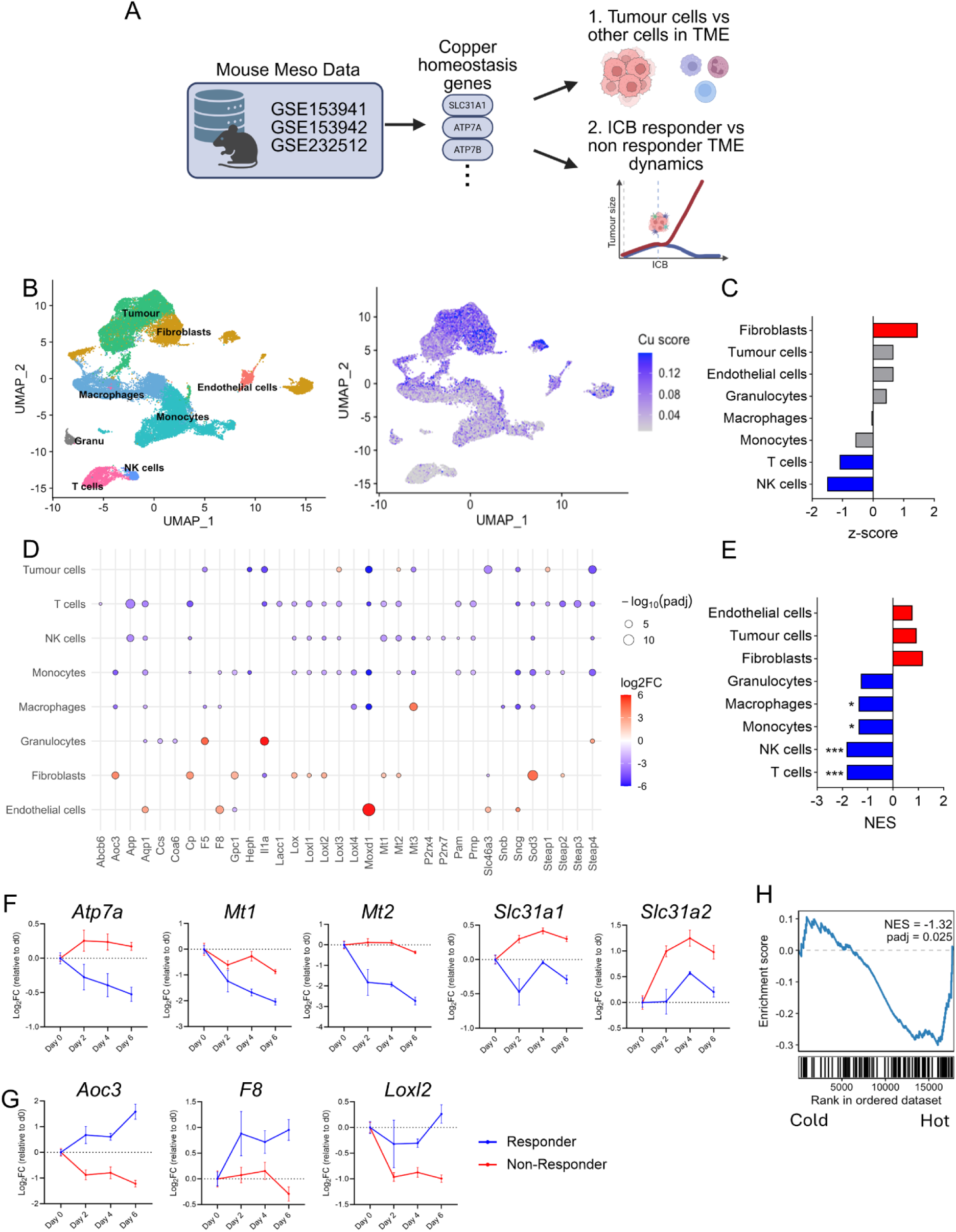
Copper transport, buffering and cuproenzyme genes are enriched in tumour cells and temporally remodelled during effective ICB. (**A**) Schematic depicting analysis of publicly available mouse mesothelioma datasets, created with BioRender.com. (**B**) UMAP projection of single-cell RNA-seq data from murine mesothelioma tumours (AB1), annotated with major cell populations (left) and overlaid with a per-cell copper homeostasis gene-set score (right). (**C**) Average expression of copper homeostasis gene set for indicated cell subsets, shown as z-scores. (**D**) Differential expressed genes (DEGs) of copper homeostasis genes across major cell types. DEGs with p-adj < 0.05 and log2FC > 1 displayed. (**E**) Normalised enrichment scores (NES) from fgsea for copper homeostasis across cell subsets. Cell subsets significantly enriched for copper homeostasis genes indicated by asterisks. (**F**, **G**) Temporal dynamics of selected copper homeostasis genes in ICB responders (R) and non-responders (NR) that had a significant response by timepoint interaction effects by DESeq2, indicating differential expression trajectories between R and NR over time. Expression is shown as log2 fold-change gene expression across three time points across post ICB timepoints (Day 2, Day 4 and Day 6), relative to pre ICB baseline (Day 0). Error bars indicate SEM. (**H**) GSEA of bulk RNA-seq data from 111 asbestos-derived mouse tumours from 72 strains of collaborative cross mice displaying enrichment of copper homeostasis gene set in high immune infiltrating compared to low immune infiltrating tumours. * padj < 0.05, ***padj < 0.005 with Benjamini–Hochberg correction for multiple comparisons.

### Copper homeostasis genes exhibit response-dependent expression dynamics during ICB treatment

Immune responses to ICB differ markedly between responders and non-responders, and we previously generated bulk transcriptomic datasets profiling responding and non-responding murine mesothelioma tumours over the course of ICB (1 hour pre-ICB/day 0, day 2, 4 and 6 after ICB) (*14*). We queried whether responding and non-responding tumours to ICB would display distinct patterns of change in copper homeostasis genes, which would inform how best to target copper metabolism in combination with ICB. Dynamic profiling identified differences in temporal expression patterns of copper homeostasis genes between responders and non-responders during ICB treatment. Specifically, we focused on genes with significant response and timepoint interaction effects, indicating that their expression trajectories differed between responders and non-responders over time (fig. S2C). Core transport and storage genes, including *Atp7a, Mt1, Mt2, Slc31a1*, and *Slc31a2*, displayed differential expression patterns over time in relation to ICB response, with non-responders maintaining or increasing expression while responders showed reduced expression over time (Fig. 2F). Conversely, vascular and extracellular matrix remodelling genes, including *Aoc3, F8* and *Loxl2*, were maintained in responders (Fig. 2G). Collectively, this data shows that effective ICB in this model is associated with distinct temporal remodelling of copper homeostasis genes, with non-responders preferentially maintaining copper transport pathways. These differing trajectories also suggest that copper metabolism is dynamically regulated during ICB and may provide a rationale for copper-targeting strategies to improve anti-tumour immunity.

Lastly, we analysed previously published bulk transcriptomic data from 111 different asbestos derived tumours from 72 different strains of Collaborative Cross mice (*15*), and found that the copper homeostasis gene set was preferentially enriched in tumours with low immune infiltration compared with tumours characterised by high expression of immune-related genes and abundant immune-cells (Fig. 2H). These data indicate that the negative relationship between copper homeostasis and immune infiltration is not restricted to a single mesothelioma model, and that these mouse models provide a tractable in vivo system to test how copper-targeting agents influence anti-mesothelioma immunity.

### TETA decreases intratumoural copper and reshapes the lymphoid compartment

As increased intratumoural copper is associated with immunosuppression and reduced immune infiltration, we hypothesised that copper chelation therapy would reduce intratumoural copper and remodel the TME. We therefore treated mice bearing subcutaneous AB1-HA or AE17-OVA tumours with the copper chelator TETA in vivo at previously optimised doses (*8*) (Fig. 3A, B). After 7 days of oral TETA, tissue copper levels were significantly decreased in the stomach, but not in the kidney, liver or lungs, consistent with the route of drug delivery (fig. S3A, B). Systemically, serum copper concentrations were unchanged, whereas urinary copper levels were significantly increased following TETA treatment (fig. S3C, D), indicating effective chelation and enhanced copper excretion. X-Ray Fluorescence microscopy (XFM) showed that copper was broadly distributed throughout untreated tumour sections but was significantly reduced following TETA treatment in both models (Fig. 3C, D). XFM analysis of additional trace elements revealed no significant differences in intratumoural calcium, iron, zinc, bromine, or sulphur between treatment groups, with bromine showing a trend toward reduction with TETA, indicating minimal off-target perturbation of other elements within the tumour (fig. S3E-I).

**Fig. 3.**
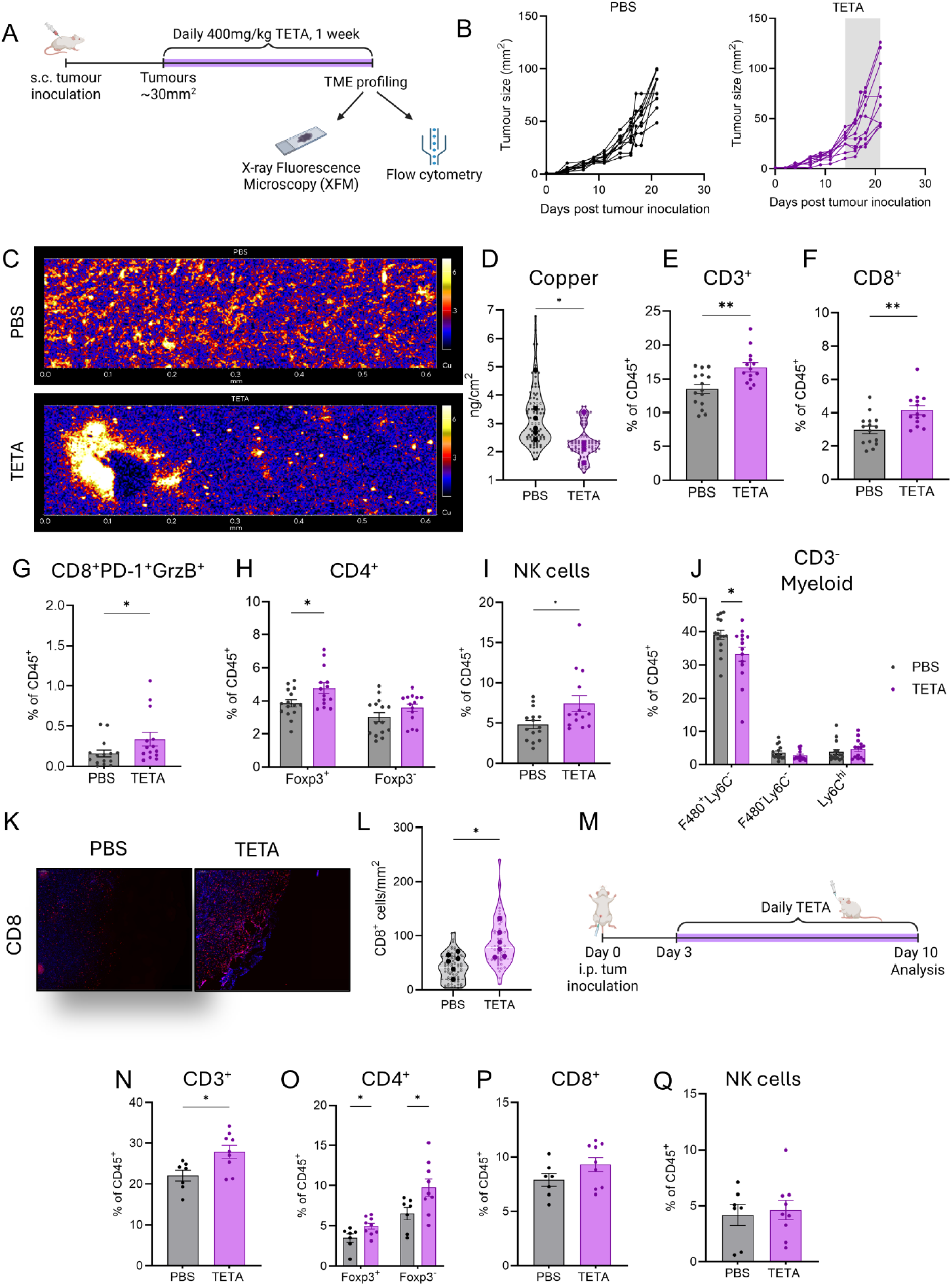
TETA reduces intratumoural copper and remodels the mesothelioma microenvironment by increasing lymphocyte infiltration. (**A**) Experimental design of 1 week TETA treatment in subcutaneous (s.c) AB1-HA or AE17-OVA tumour bearing mice, created with BioRender.com. (**B**) Representative tumour growth curves of subcutaneous AB1-HA tumours in vehicle (PBS) versus TETA treated animals. TETA treatment duration in grey. (**C**) Representative XFM elemental map displaying spatial copper concentration by colour intensity in PBS and TETA treated s.c tumour sections. (**D**) Tumoural copper concentrations (ng/cm^2^) in PBS and TETA treated tumours. Violin plots display the distribution from 90 individual measurements from 15 regions per treatment (n = 6/group), nested within individual biological samples in dots. Dot plots representing AB1-HA s.c tumour infiltrating (**E**) CD3^+^, (**F**) CD8^+^, (**G**) CD8^+^PD-1^+^GrzB^+^, (**H**) CD4^+^Foxp3^+/-^ T cells, (**I**) CD335^+^ NK cells and (**J**) F480^+/-^Ly6C^hi/-^myeloid cells, as a percentage of CD45^+^ cells, TETA versus PBS (n=14-15/group). (**K**) Representative immunofluorescence images of CD8^+^ T cells (red), and DAPI nuclei (blue) in s.c AE17-OVA tumours from PBS versus TETA treated animals. (**L**) Violin plots display CD8^+^ cells/mm^2^ quantification, distribution of 70 individual measurements per treatment nested within individual biological samples (n = 6/group). (**M**) Experimental design of 1 week TETA treatment in intraperitoneal (i.p) AB1-HA tumour bearing mice, created with BioRender.com. Dot plots representing AB1-HA i.p tumour infiltrating (**N**) CD3^+^, (**O**) CD4^+^Foxp3^+/-^, (**P**) CD8^+^ T cells and (**Q**) CD335^+^ NK cells as a percentage of CD45^+^ cells (n = 7-9/group). All flow cytometry data presented as mean ± SEM, with two-tailed Mann-Whitney test used to compare differences between treatment groups. Nested t-test used to compare differences between treatment groups for XFM and immunofluorescence data. *p < 0.05, **p < 0.01.

We further characterised how TETA remodels the immune compartment by flow cytometry analysis in AB1-HA tumours. CD45⁺ cell frequencies and absolute numbers were similar between TETA and control tumours (fig. S4A, B). TETA treatment increased the proportion of CD3^+^ T cells (Fig. 3E) and CD8^+^ T cells (Fig. 3F). The overall frequencies of CD8^+^ T cells expressing inhibitory or activation receptors KLRG-1, LAG3, PD-1, PD-L1 were similar between TETA and vehicle treated mice (fig. S4C). However, the proportion of CD8⁺ T cells co-expressing granzyme B and PD-1 was increased following TETA treatment (Fig. 3G), suggesting an increase in activated, cytotoxic CD8⁺ T cells within the tumour. Among CD4⁺ T cells, the frequency of CD4⁺Foxp3⁺ regulatory T cells (Tregs) increased with TETA, whereas CD4⁺Foxp3⁻ T cells (Tconv) were unchanged (Fig. 3H). The expression of inhibitory or activation markers on both Tconv and Tregs was comparable between treatment and control groups (fig. S4D, E). An increased proportion of NK cells was observed with TETA treatment (Fig. 3I). Within the myeloid compartment, the frequency of CD3⁻F4/80⁺Ly6C⁻ macrophages was reduced in TETA treated tumours (Fig. 3J), plasmacytoid dendritic cell (pDCs) and conventional dendritic cells (cDCs1/2) proportions were similar between groups (fig. S4F). We assessed tumour draining lymph nodes (DLN) and found similar proportions of CD4^+^ Tconv, Tregs, naïve, effector, memory CD8^+^ T cells, NK cells and DC populations between treatment and control groups (fig. S4G-K). Tumour antigen cross-presentation, assessed by proliferation of transferred HA-specific TCR-transgenic CD8⁺ T cells, was comparable between groups (fig. S4L), consistent with TETA exerting its effects within the tumour rather than the DLN.

To test whether TETA driven remodelling of the tumour lymphoid compartment was observed in different mesothelioma models, we assessed the same TETA treatment in subcutaneous AE17-OVA tumours on a different genetic background (C57Bl/6). IHC indeed showed increased CD8⁺ infiltration in TETA-treated tumours compared with controls (Fig. 3K, L). We also used an intraperitoneal AB1 model (Fig. 3M) and observed that TETA increased the proportion of CD3⁺ T cells, driven primarily by CD4⁺ T cells and less so CD8^+^ T cells, while NK cell proportions were unchanged (Fig. 3N-Q). Together, these data show that TETA selectively remodels the subcutaneous TME toward an inflammatory, ICB-responsive state, including increased intratumoural CD8⁺ T cells with enhanced cytotoxic activation and reduced macrophage abundance, while intraperitoneal tumours show a distinct pattern of T-cell remodelling.

### TETA remodels inflammatory gene programs in the TME

To define the transcriptional changes resulting from copper depletion within the TME, we performed bulk and single-cell transcriptomic profiling of subcutaneous AB1-HA tumours from TETA treated and control mice (Fig. 4A). Bulk RNA sequencing showed that TETA treated tumours were enriched for a pro-inflammatory gene signature in murine tumours that were responsive to ICB treatment (*16*), whereas control tumours were enriched for the copper homeostasis gene set (Fig. 4B). We further interrogated TETA mediated changes in the tumour microenvironment by single-cell RNA sequencing. After quality control, dimensionality reduction and clustering we obtained 47,352 cells from the malignant and stromal components (tumour, mesenchymal stromal-like cells), and 14,048 immune cells, including macrophages, monocytic derived macrophages, CD8^+^ T cells, NK cells, Tregs, γδ T cells (Tgd), neutrophils, B cells, dendritic cells, mast cells and basophils (Fig. 4C, fig. S5A). The number and proportion of cell clusters were broadly comparable between TETA and control groups (Fig. 4D, E).

**Fig. 4.**
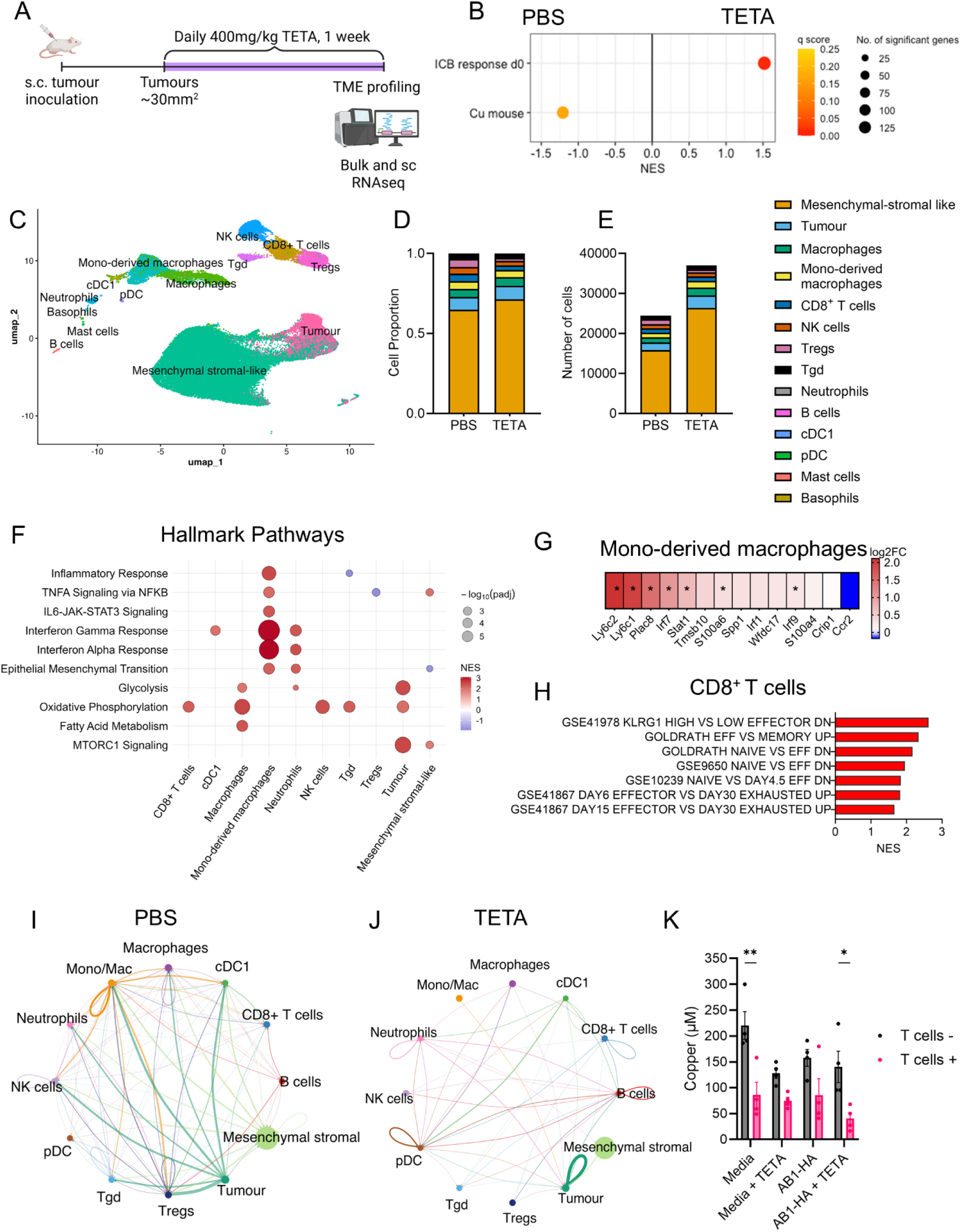
TETA drives a proinflammatory tumour myeloid gene signature and effector CD8^+^ T cell programming. (**A**) Experimental design of 1 week TETA treatment in subcutaneous (s.c) AB1-HA tumour bearing mice, created with BioRender.com. (**B**) GSEA plots displaying ICB response and copper homeostasis gene sets significantly enriched (q < 0.25) in AB1-HA tumours from PBS or TETA treated animals. (**C**) UMAP projection single-cell RNA-seq data AB1-HA tumours, with (**D**) proportions and (**E**) numbers of cells from each cluster, split by treatment. (**F**) Bubble plot depicting GSEA plot for HALLMARK pathways using the fgsea package. Significantly enriched pathways in TETA-treated (red) or PBS-treated (blue) (padj < 0.05) across individual cell clusters. (**G**) Heatmap of log_2_FC in expression between treatment arms for genes associated with proinflammatory monocytes, significantly enriched individual genes indicated with * (padj< 0.05). (**H**) Bar graphs of significantly enriched (padj < 0.05) IMMsigDB T cell effector and exhaustion pathways in CD8^+^ T cells from TETA treated samples. Circos plots from CellChat analysis showing interactions with inferred communication strength in (**I**) Control relative to TETA, and (**J**) in TETA relative to Control (right, TETA up). Node size reflects cell abundance and edge width reflects the relative increase in communication strength for each interaction. (**K**) Concentration of copper in AB1-HA +/- TETA conditioned media after 30 minutes incubation with mouse CD8^+^ T cells isolated from BALB/c splenocytes. Data presented as mean ± SEM, (n = 4/condition), biological replicates, two independent experiments. Two-tailed paired t-test used to compare differences between T cell and no T cell controls.

Differential expression analysis revealed that the transcriptional response to TETA was highly cell type specific, with the largest number of Differentially Expressed Genes (DEGs) in macrophages and monocytic-derived macrophages (fig. S5B). TETA treatment was associated with enrichment of interferon and inflammatory response related pathways in monocytic derived macrophages, alongside oxidative phosphorylation pathways in CD8+ T cells and NK cells (Fig. 4F, fig. S5C). We previously identified an IFN-producing inflammatory monocyte population as a key determinant of ICB response in our model (*14*). Analysis of monocytic-derived macrophages showed enrichment of this pro-inflammatory monocyte signature following TETA, alongside a trend towards an increased proportion of these cells in the dataset (Fig. 5G, fig. S5D). Effector-like gene programs were also enriched in CD8^+^ T cells from TETA-treated samples. (Fig. 5H). Inferred cell–cell communication analysis revealed substantial remodelling of the inferred signalling network following TETA treatment (Fig. 4I, J), as TETA reduced tumour-derived interactions directed towards lymphoid and myeloid populations that involved extracellular matrix (ECM) remodelling, collagen, fibronectin 1 (FN1) and Secreted Phosphoprotein 1 (SPP1) and amyloid precursor protein (APP) pathways (fig. S5E). We next assessed whether TETA alters copper availability to T cells using a staggered co-culture system with AB1-HA tumour conditioned media. AB1-HA conditioned media supported minimal T cell copper uptake, but AB1-HA and TETA-conditioned media promoted greater copper uptake by T cells (Fig. 4K). Together these findings suggest that TETA shifts the TME to a more proinflammatory landscape that favours anti-tumour immunity.

**Fig. 5.**
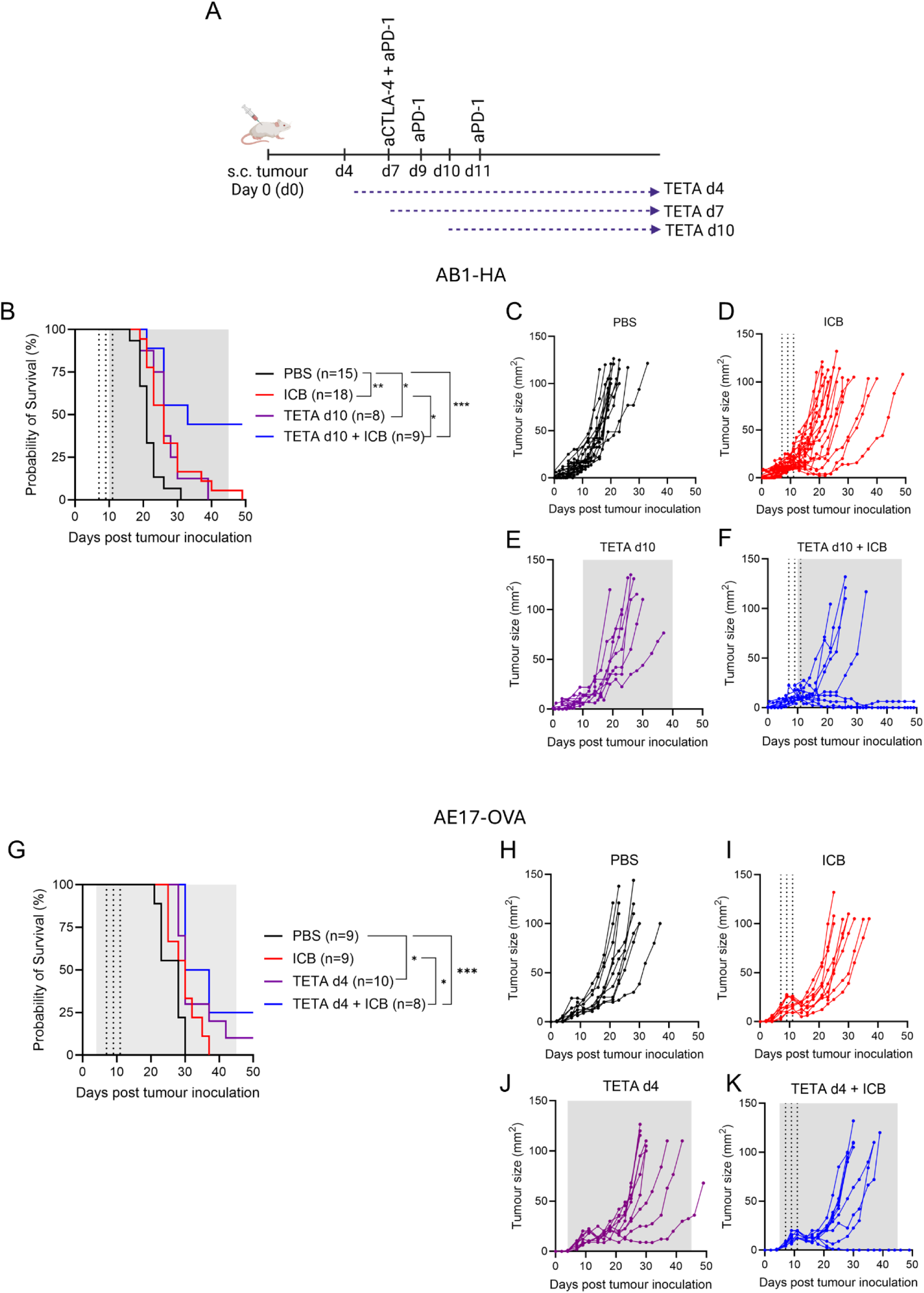
TETA in combination with ICB improves median overall survival and tumour control in a subset of mice. (**A**) Experimental timeline for different TETA dosing schedules. (**B**) Kaplan-Meier survival curves of subcutaneous AB1-HA tumours in mice treated with vehicle control (PBS, black), immune checkpoint blockade (ICB; anti-CTLA-4 and anti-PD-1; red), TETA (purple) and combination TETA and ICB (blue). TETA initiated at day 10 (d10) post tumour inoculation. Individual tumour growth curves for (**C**) PBS, (**D**) ICB, (**E**) TETA, (**F**) TETA + ICB. (**G**) Kaplan–Meier survival curves of subcutaneous AE17-OVA tumours in mice treated with PBS, ICB, TETA and combination TETA and ICB. For AE17-OVA, TETA was initiated at d4 after tumour inoculation. Individual tumour growth curves for (**H**) PBS, (**I**) ICB, **J**) TETA, (**K**) TETA + ICB treatment. Pairwise comparisons for survival were performed using a Mantel–Cox log-rank test (*p < 0.05, ***p < 0.005). Dotted vertical lines denote ICB treatment time points with grey shaded areas indicating TETA treatment. Group sizes indicated in figures.

### TETA enhances dual CTLA-4/PD-1 ICB responses in a schedule-dependent manner

Given that TETA shifted the TME to a more inflamed, cytotoxic state, we next determined whether adding TETA to dual anti-CTLA-4 and anti-PD1 ICB could enhance anti-tumour immunity. As copper homeostasis genes in ICB responders and non-responders exhibited time-dependent dynamics, we varied the timing of TETA administration relative to ICB: before ICB (day 4; d4), concurrently with ICB (day 7; d7) and after initiation of ICB (day 10; d10) (Fig. 5A). In the AB1-HA model, both ICB alone and TETA given at d10 delayed tumour growth and improved overall survival compared to PBS alone (Fig. 5B-F). Notably, combining TETA at d10 with ICB further improved overall survival and increased the number of complete responders relative to ICB alone (Fig. 5B, F). Administering TETA before (d4) or concurrently with (d7) ICB also yielded some long-term responders, but median overall survival was not significantly different from ICB alone (fig. S6A–F). In a second mesothelioma model (AE17-OVA) that is highly resistant to the same ICB regimen, TETA given before ICB (d4) produced the greatest benefit, significantly increasing median overall survival and the number of long-term survivors compared with ICB alone (Fig. 5G–K). In contrast, administering TETA concurrently with or after ICB in this resistant model did not improve median overall survival (fig. S6G–L). Overall, these studies show that copper chelation with TETA can enhance ICB efficacy in mesothelioma, likely by rendering the TME more responsive to immune activation.

## DISCUSSION

Here we examined the role of copper in shaping the TME and anti-tumour immunity. In murine mesothelioma models, copper depletion with a chelator (TETA) decreased intratumoural copper, and reprogramed the TME by increasing expression of proinflammatory genes and immune cell infiltration. In preclinical models of mesothelioma, TETA improved anti-tumour immunity in combination with anti-CTLA-4 and anti-PD-1 ICB. TETA is a clinically approved copper chelator with an established safety profile from long-term use in Wilson’s disease, making it an attractive candidate for therapeutic repurposing. These findings establish the rationale for using copper chelation as a strategy to potentiate the effects of ICB.

In both human and mouse mesothelioma, copper homeostasis genes compartmentalised to malignant cells and fibroblasts, whereas lymphocyte populations displayed downregulation of this program. Consistent with this pattern, mouse mesothelioma tumours with low immune infiltration were enriched for copper homeostasis genes. Although copper accumulation in tumour tissue has been reported previously (*10*), our study suggests that this accumulation is not uniform across the TME. In support of this, recent spatial mass spectrometry analysis of murine 4T1 breast tumours showed copper accumulation in regions with minimal immune infiltrate, raising the possibility that tumour-infiltrating immune cells reside in relatively copper-limited niches (*17*). Building on this spatial perspective, our data support a model in which copper is not only heterogeneously distributed but also differentially available across cellular compartments within the tumour microenvironment. Malignant and stromal cells, which exhibit high expression of copper transport and buffering machinery, may sequester bioavailable copper and limit copper availability to infiltrating immune cells. In this context, altered copper availability to tumour-infiltrating T cells may constrain their activation and cytotoxic function, given the requirement for copper in processes such as mitochondrial respiration and effector activity (*18*). Our findings raise the possibility that TETA may modify this imbalance by redistributing copper within the tumour microenvironment. Although the precise in vivo mechanism remains to be demonstrated, TETA may act beyond simple copper depletion by mobilising tumour-associated copper pools and increasing extracellular copper bioavailability. Consistent with this proposed mechanism, conditioned media from TETA treated tumour cells supported increased copper uptake by CD8⁺ T cells, suggesting that tumour cell exposure to TETA can influence copper availability to lymphocytes. In this way, TETA may alleviate tumour associated metabolic constraints and create conditions more permissive for effective anti-tumour T cell responses. These observations underscore the complexity of copper biology in mesothelioma and suggest that modulating intratumoural copper may have broad consequences across malignant, stromal, and immune compartments.

Our study revealed that copper-homeostasis genes were dynamically regulated in association with ICB responses, with distinct time-resolved patterns. Because these highly controlled models showed no difference in tumour size between responders and non-responders at the time of sequencing (*14*), the observed transcriptional differences are unlikely to simply reflect tumour burden and instead point to rewiring the TME. A limitation is that these analyses were derived from bulk RNA sequencing and therefore could not resolve cell type specific dynamics. Although copper-related genes have previously been linked to survival (*12, 19*) and proposed as biomarkers of therapeutic response (*20, 21*), our findings suggest that these gene programs are dynamic, and that the timing of measurement may be critical for biomarker development.

The key implication of our study is that TETA remodelled the TME toward an ICB responsive inflammatory state, characterised by increased intratumoural cytotoxic lymphocyte features and enrichment of inflammatory gene programs. This did not reflect a global increase in leukocyte infiltration, as total CD45^+^ cell numbers were unchanged, and TETA also increased the proportion of CD4^+^ Tregs, pointing to a shift in immune composition within the tumour. Similar increases in tumour infiltrating lymphocytes have been reported in other tumour models with different in vivo copper chelators and treatment durations (*6, 8, 9, 19*). Although expression of activation and inhibitory markers on CD8^+^ T cells was broadly similar between groups, single-cell analysis suggested enrichment of genes associated with oxidative phosphorylation (OXPHOS) and effector programming in CD8^+^ T cells with TETA. Enhanced OXPHOS has been linked to improved CD8^+^ T cell infiltration into tumours (*22*), and copper has recently been shown to support OXPHOS-dependent T cell function in autoimmunity (*23*), raising the possibility that altered copper availability rewires lymphocyte metabolism. In the absence of major shifts in surface phenotype, these data suggest that TETA induces fine-tuning of the lymphocyte state. Consistent with this, these effects were confined to the tumour and not in DLNs, supporting a local effect of TETA within the TME.

In line with lymphocyte fine-tuning, TETA enhanced dual CTLA-4/PD-1 blockade in mesothelioma models. Although a prior study with the copper chelator Tetraethylenepentamine (TEPA, Cuprior analogue) showed reduced tumour growth when combined with anti–PD-1 therapy in murine gastric cancer (*19*), to our knowledge this is the first study to demonstrate that TETA can improve long-term survival in combination with ICB, including complete tumour regressions in a subset of mice. Durable benefit has also been reported with a copper-depleting nano-agent composed of bacterial vesicles loaded with a copper chelator in combination with anti–PD-1 therapy in breast cancer models, although the contribution of the immunogenic vesicle platform itself was not defined (*17*). Given apparent fine-tuning of lymphocytes in the TME by TETA, it will be important to determine whether TETA can enhance other T cell based immunotherapies, such as CAR T cells or TIL therapy. The model dependent difference in optimal timing of TETA relative to dual ICB could reflect the kinetics of ICB-induced TME changes. In the AB1 model, copper transport (*Slc31a1, Atp7a, Mt1*) genes were sustained post-ICB in non-responders, consistent with the finding that TETA was most effective when administered after ICB was initiated in the AB1-HA model. Although ICB-response dynamics were not assessed in AE17, tumour-infiltrating CD8^+^ T cells in AE17 displayed more activated/exhausted like phenotypes (*24*). Earlier copper modulation could be required to prime the AE17 TME by increasing CD8^+^ T cells that are more responsive to ICB. Despite the differences in timing of initiation, both models indicate that sustained copper chelation alongside ICB drives robust anti-tumour immunity, providing a rational to assess TETA either prior to, or during the first few cycles of ICB clinically. Due to the known variability of ICB response kinetics between patients and tumour types, monitoring tumour or circulating metal levels may provide a pharmacodynamic approach to guide the timing of when to best initiate copper-targeted therapy.

Although we previously reported that TETA altered tumour-infiltrating neutrophil phenotype in a neuroblastoma cancer model (*8*), changes in our mesothelioma model involved macrophages, which showed the largest transcriptional responses to treatment by single-cell analysis. Consistent with prior flow cytometry studies, TETA reduced overall tumour-infiltrating macrophages (*19, 25*). However, our single cell data also revealed that TETA reshaped the macrophage compartment, enriching pro-inflammatory transcriptional programs in monocytic-like macrophages, including a monocyte signature previously associated with ICB response (*14*). CellChat analysis further suggested that TETA broadly disrupted tumour-derived signalling to immune populations, supporting the idea that copper chelation reshapes not only immune cell composition but also the communication networks that sustain the local suppressive microenvironment. Although our study focused primarily on lymphoid remodelling because of its relevance to ICB response, these data suggest that myeloid reprogramming and altered stromal–immune signalling may contribute to the overall TETA response.

Our findings show that TETA depletes intratumoural copper, reshapes the mesothelioma TME, and enhances dual immune checkpoint blockade, providing a strong translational rationale for clinical testing. These data support evaluation of TETA in combination with first-line ICB for mesothelioma.

## MATERIALS AND METHODS

### Mice

All experiments used 8 to 12-week-old male and female BALB/c and C57BL/6 mice obtained from the Ozgene Animal Resource Centre (Murdoch, Australia) or the Harry Perkins Institute of Medical Research (Nedlands, Australia). Clone 4 (CL4xThy1.1) T cell receptor (TCR) transgenic mice express a TCR that recognises a major histocompatibility complex (MHC) class I-restricted influenza A/PR/8 hemagglutinin (HA_533−541_) epitope. Mice were housed at the Harry Perkins Institute of Medical Research Bioresources Facility (Nedlands, Australia) under standard specific pathogen-free housing conditions. All animal experiments were conducted under the approval of the Harry Perkins Institute of Medical Research Animal Ethics Committee (protocol AE271, 384).

### Tumour cell lines

Murine mesothelioma cell lines were derived as previously described (*26–28*). AB1 was transfected with the model antigen-HA (haemagglutinin), and AE17 transfected with sOVA (secretory ovalbumin) to enable tracking of model tumour antigen-specific responses, or identification of tumour cells in single cell sequencing experiments where described. Cell lines were maintained in RPMI 1640 (ThermoFisher Scientific) supplemented with 20 mM HEPES (ThermoFisher Scientific), 0.05 mM 2-mercaptoethanol, 100 units/mL penicillin (CSL, Melbourne, Australia) and 10% newborn calf serum (NCS; ThermoFisher Scientific). Cell lines were kept under geneticin (500 µg/mL G418; ThermoFisher Scientific) selection pressure to select for HA and OVA expressing cells. All cell lines were tested for Mycoplasma spp. every four months and remained negative. Tumour cells were harvested at 80% confluence after a minimum of 4 passages after thawing (maximum of 10 passages). Mice were inoculated subcutaneously in the right-hand flank or intraperitoneally with 5 x 10^5^ tumour cells where described.

### Copper chelation and ICB treatments

Triethylenetetramine (TETA, Sigma-Aldrich) was administered by oral gavage at a dose of 400 mg/kg/day (*8*) for 7 days in characterisation experiments. In combination therapy experiments, TETA was administered at 800mg/kg daily for 14 days, followed by once every alternate day up to a total of 25 doses. ICB consisted of anti-CTLA-4 (4F10) dosed once at 100 μg/mouse and anti-PD-1 (RMP1-14) dosed three times at 200 μg/mouse with two-day intervals as previously described (*29*). Responders to therapy were defined as complete tumour regression to 0 mm^2^ that was maintained for a minimum of 30 consecutive days. Mice were randomised to treatment groups. Investigators measuring tumour sizes and performing downstream analysis were blinded to treatment groups.

### Atomic absorption spectrophotometry

Serum, urine and organs were collected. Wet weights of organs were recorded, and organs were freeze-dried overnight. Copper measurements by atomic absorption spectrophotometry were performed as previously described (*6*). Dry organ weights were recorded, digested in 200 μL of hydrogen peroxide and 250 μL of nitric acid, heated for 2 hours at 80°C and resuspended with 4 mL of MilliQ water. Samples were diluted and analysed on the graphite furnace component of the PinAAcle 900T dual flame/graphite furnace atomic absorption spectrophotometer (PerkinElmer, Australia) using Zeeman background correction. All measurements were performed in triplicate.

### Flow cytometry

Freshly harvested tumours and tumour-draining axillary and inguinal lymph nodes (DLNs) were prepared to single-cell suspensions for flow cytometry as previously described (*24*). Briefly, tumours were minced into roughly 2 mm^2^ sized pieces and enzymatically digested with 1.5 mg/mL type IV collagenase (Worthington Biochemical) and 0.1 mg/mL deoxyribonuclease (DNAse, Sigma Aldrich) in PBS + 2% NCS at 37°C for 1 hour. DLNs were enzymatically digested with 1 mg/mL type IV collagenase (Worthington Biochemical) and 1 μg/mL DNase (Sigma Aldrich) in RPMI1640 supplemented with 2% NCS and 20 mM HEPES (ThermoFisher Scientific) for 25 minutes at room temperature. Ethylenediaminetetraacetic acid (EDTA; Sigma Aldrich) was added during the last five minutes of digestion, and the resulting cell suspension was filtered using a 40 μm strainer and underlaid with 2 mL of cold 0.1 M EDTA-NCS (1:10) before centrifugation without brake to enrich for viable cells for downstream analyses. The spectral flow cytometry panel used to characterise lymphoid and myeloid cell subsets is outlined (table S3). Briefly 1 x 10^6^ cells from each sample were stained with the Fc receptor block anti-CD16/CD32 for 30 minutes at 4°C. To identify live cells, samples were stained with a viability stain, ViaDye Red, diluted in PBS + 1 mM EDTA and incubated at 4°C for 20 minutes. All surface staining antibodies were diluted with PBS + 2% NCS and incubated at 4°C for 30 minutes. For intracellular staining, cells were permeabilised using the Foxp3/Transcription Factor Staining Buffer Set (ThermoFisher), washed with Permeabilization Buffer (ThermoFisher) and stained. All antibodies for intracellular staining were diluted in Permeabilization Buffer (ThermoFisher) and incubated at 4°C for 30 minutes. Single stain and fluorescence minus-one (FMO) staining were also performed. Data was acquired on the Cytek® Aurora (Cytek Biosciences, California, USA) with 100,000 live events collected per sample where possible. All flow cytometry analyses were conducted using FlowJo™ Software version 10.10.0 (BD Biosciences, New Jersey, USA). A summary of the gating strategy used is outlined (fig. S7).

### Bulk RNA sequencing of murine tumour samples

Whole tumours were harvested and stored in RNAlater (Life Technologies) at -80°C. RNA was extracted from frozen tumours using the RNeasy Plus Mini Kit and Tissue Ruptor (QIAGEN). RNA quality was confirmed using a Bioanalyzer (Agilent Technologies). Library preparation and sequencing (2x150-base pair paired end using an Illumina Novaseq6000 platform) was performed at the Medical Genomics Laboratories (Murdoch, WA, Australia).

### Analysis of publicly available mesothelioma gene expression data

Mesothelioma tumour gene expression and clinical data from The Cancer Genome Atlas Program (TCGA-MESO, n = 87) were retrieved using TCGAbiolinks (*30*). A second dataset (GSE51024) included pleural mesothelioma tumours (n = 55) and matched normal lung parenchyma samples (n = 41) (*11*). Expression data were harmonised in a common analytical framework and log2-transformed where required before downstream analyses. For cross-cohort comparisons, datasets were aligned by shared gene identifiers and adjusted for dataset-of-origin batch effects using ComBat with empirical Bayes shrinkage (*31*). Batch was defined by dataset source, and biological covariates of interest, including sample type and histological subtype where applicable, were retained in the model. Genes with zero variance across samples were excluded from batch correction and retained uncorrected in the final expression matrix. Expression was compared between lung and tumour samples. Overall survival was assessed in tumour samples using Kaplan-Meier and Cox proportional hazards analyses. Gene set activity was quantified using GSVA with the ssGSEA method where applicable.

Gene expression data from mouse mesothelioma tumour samples collected before ICB (day 0, d0) and at days 2, 4, and 6 (d2, d4, d6) following ICB were obtained from GSE153941, with tumours classified as responders or non-responders according to the original study (*14*). Temporal expression changes in relation to ICB response were modelled for all genes using DESeq2 with : *∼ response + timepoint + response:timepoint*, with day 0 as the reference timepoint. Copper homeostasis genes with a significant response-by-timepoint interaction at any post baseline timepoint were selected for visualisation. For each gene, expression was expressed as log2 fold change relative to Day 0 mean and plotted over time for responders and non-responders.

Gene expression data from 111 asbestos-induced primary mouse mesothelioma samples in 72 genetically diverse Collaborative Cross mouse strains were obtained from GSE232512. Samples were classified as immune or non-immune according to the original study (*15*). GSEA was performed on normalised gene expression data for a copper homeostasis gene set. Enrichment with an FDR q value < 0.25 was considered significant.

### X-Ray Fluorescence Microscopy (XFM)

Metal distribution maps were acquired at the Australian Synchrotron X-ray fluorescence microscopy (XFM) beamline (*32, 33*). Tumour cryosections (7 µm thick) were mounted on 1 µm thick silicon nitride windows (Melbourne Centre for Nanofabrication), air dried, and analysed under ambient conditions. A Si 111 double multilayer monochromator was used to tune a15.8 keV X-ray beam, focused using a Kirkpatrick–Baez mirror pair to an approximately 2 µm full width at half maximum spot size. Elemental emission spectra were collected in event-mode using a Vortex ME3 detector array with three active detector elements, at a 90° collection angle geometry relative to incident beam. Initial overview scans were performed to localise tumour regions. Quantitative copper maps were acquired by raster scanning with a 2 µm step size and 2.5 ms dwell time per pixel. For selected regions, higher-resolution scans were performed at 1 µm step size with equivalent dwell time. A tissue matrix model (7 µm thickness, 1.0 g/cm³ density) incorporating a 100 nm silicon nitride support layer was used for absorption correction and concentration estimation.

Elemental quantification was performed on XFM maps processed using GeoPIXE (ver.7.7f, CSIRO, Australia). For each tumour section, fifteen non-overlapping regions of interest (ROIs) were manually defined. Each ROI comprised 1,638 pixels (∼ 6,552 µm²; pixel size 2 µm × 2 µm). ROIs were selected based on tissue density and morphology to identify viable tumour regions and were placed independently of elemental signal intensity to minimise selection bias. To capture spatial heterogeneity, five ROIs were positioned at the tumour periphery, five in the mid-interior, and five in the central region. Elemental concentrations for bromine (Br), calcium (Ca), copper (Cu), iron (Fe), sulphur (S), and zinc (Zn) were initially quantified in parts per million (ppm). Areal densities (ng/cm²) were calculated from the tissue matrix model assumptions using the following formula: ng/cm^2^ = ppm × 0.7. ROIs were treated as technical replicates nested within individual tumours, while tumours were treated as independent biological replicates.

### Immunofluorescence

Formalin-fixed and paraffin-embedded (FFPE) tumour sections (5 µm) were deparaffinised, rehydrated, and subjected to heat-induced antigen retrieval. Endogenous peroxidase activity was quenched using Peroxidazed-1 (Biocare, USA), and sections were blocked with Background Sniper (Akoya Biosciences, USA). Sections were stained with rabbit anti-mouse CD8α antibody (Cell Signaling Technology, USA; clone D4W2Z; #98941), followed by a poly-HRP anti-rabbit secondary reagent (Thermo Fisher SuperBoost™ Goat anti-Rabbit Poly HRP, USA; #B40962) and Alexa Fluor 594 tyramide amplification (Thermo Fisher, USA; #B40957). Sections were imaged on a Panoramic 250 FLASH III Digital Scanner and exported as TIF files using SlideViewer (3D Histech, Hungary) for downstream analysis.

CD8^+^ T cells were quantified in Fiji/ImageJ using Trainable Weka Segmentation with a supervised pixel-classification workflow (*34*). A classifier trained on 12 representative images (6 AB1-HA tumours and 6 AE17-OVA tumours) used annotated CD8 positive cells as the positive class, and auto-fluorescent or background regions as the negative class. Features included Gaussian blur, Hessian, membrane projections, Sobel, and difference of Gaussians (sigma 0.7-2.5). The classifier was applied in batch in Fiji/ImageJ to generate segmentation masks, and objects were quantified by particle analysis (20-500 px²; circularity 0.1-1.0). Tissue area was calculated using a fixed calibration (0.2 mm = 145.5 pixels), and CD8^+^ T cell density was reported as cells/mm².

### Single cell RNA sequencing

Tumours were dissociated as above, and stained with a viability dye (Viadye Red, 1:1000, Cytek Biosciences), CD335-APC (clone 29A1.4, 1:200, Biolegend), CD11b-SparkYG593 (clone M1/70, 1:400, BioLegend), and CD3-FITC (clone 17A2, 1:100, Cytek Biosciences). CD3^+^, CD11b^+^ and CD335^+^ populations were sorted as immune cells, and marker negative cells were sorted as tumour/stromal cells on the BD Rhapsody system (BD Biosciences, USA), targeting approximately 20,000 cells per group for transcriptomic profiling. Whole transcriptome libraries were constructed following the BD Rhapsody single-cell whole transcriptome analysis (WTA) workflow according to the manufacturer’s instructions. Libraries were quantified using a High Sensitivity DNA chip on Agilent Tapestation 2200 and the Quantus High Sensitivity double-stranded DNA Assay Kit. DNA libraries were sequenced on an Illumina NovaSeq 6000 S1 2 × 150 bp kit to yield an average of 40,000-50,000 reads per cell. FASTQ files for samples were processed on the Seven Bridges Platform using the BD Rhapsody Sequence Analysis Pipeline (Revision 18). Reads were aligned to the Mouse (mm10) Whole Transcriptome Analysis (WTA) reference (RhapRef_Mouse_WTA_2025-03). The pipeline performed quality control, gene alignment, and generated UMI cell-by-gene expression matrices.

Because tumour cells in the AB1-HA model expressed HA, reads were aligned to a custom reference containing the HA sequence to distinguish tumour cells in the single-cell RNA-seq dataset. A tumour-enriched gene signature was derived from HA-positive cells and then scored across all cells using Seurat. Cells with tumour-signature scores above the 90^th^ percentile and low or absent *Ptprc* (CD45) expression were classified as tumour cells.

Downstream single cell analysis was conducted in Seurat (v5.0). Low-quality cells with more than 10% mitochondrial content or fewer than 100 detected features were excluded, leaving 64,451 cells. Data from multiple samples were then normalised and integrated in Seurat using SCTransform, followed by Principal Component Analysis (PCA) and Uniform Manifold Approximation and Projection (UMAP). Cell-cycle phases were determined using Seurat’s CellCycleScoring function with the help of cc genes. Clustering was performed using Louvain and Leiden clustering in Seurat v5. Cell clusters were initially annotated using the ImmGen reference dataset for mouse via the SingleR package and further curated manually based on cluster-enriched marker genes identified by differential expression analysis and inspection of average gene expression after batch correction.

### Publicly available mesothelioma single-cell RNA-seq datasets

Single-cell RNA-sequencing datasets from human mesothelioma samples (GSE190597; EGAD50000001706) and mouse mesothelioma tumour samples (GSE153942) were obtained from publicly available repositories. Where available, the original cell type annotations provided by the respective studies were retained. Processed Seurat objects were used as the starting point for downstream analyses.

### Single cell RNA sequencing analysis

Copper gene activity was scored per cell using AddModuleScore on a curated copper homeostasis gene list, generating a metadata module score used for downstream visualisation and comparisons between cell types and groups.

Within each cell type, differential expression between condition was conducted in Seurat using the MAST framework, with relevant covariates included as latent variables to account for confounding effects. Genes were ranked by the direction and magnitude of the differential expression statistic to generate preranked lists for enrichment testing. Gene Set Enrichment Analysis (GSEA) was then performed using fast GSEA (fgsea) against the MSigDB Hallmark gene sets, and statistical significance was assessed by permutation testing. In parallel, over-representation analysis (ORA) was conducted using clusterProfiler to identify enriched Gene Ontology (GO) terms using a two-tailed Fisher’s exact test on significantly differentially expressed genes. For all enrichment analyses, p-values were corrected for multiple comparisons using the Benjamini–Hochberg procedure, and adjusted p-values were used to define significance and for downstream visualisation.

Inference of cell-cell communication networks were performed using CellChat (v2) with CellChatDB (v2). To reduce interactions driven by sparse expression, a 25% truncated mean was applied such that, within a given cell group, the average expression of a ligand–receptor component was set to zero when the proportion of expressing cells was ≤ 25%. Cell–cell communication probabilities (interaction weights) were inferred from filtered expression profiles, and signalling networks were summarised and visualised to identify dominant communication pathways across cell types and to compare condition-specific differences.

### Staggered co-culture copper transfer assay

Media alone or containing 1 × 10^6^ AB1-HA tumour cells were seeded in 10% FCS/RPMI before overnight treatment with 2 mM TETA. Tumour cells underwent centrifugation, and the resulting cell-free conditioned media was collected for co-culture with CD8^+^ T cells. Splenocytes from BALB/c mice underwent magnetic enrichment of CD8+ T cells (Miltenyi Biotec, 130-104-075) as per manufacturer’s instructions. 2 × 10^5^ CD8^+^ T cells were resuspended in respective conditioned media for 30 minutes at room temperature before centrifugation. Cell-free conditioned media (+/- T cells) were transferred to fresh tubes for copper concentration measurements.

The concentration of copper in media samples was quantitatively determined using a commercial Copper Assay Kit (Abcam, #ab272528) according to the manufacturer’s instructions. Absorbance was determined using a Benchmark Plus Plate Reader with Microplate Manager v5.2.1 (Bio-Rad, USA) at a wavelength of 359 nm.

### Statistical analysis

All statistical analysis was performed using GraphPad Prism version 10.2.3 (GraphPad Software, Massachusetts, USA). Statistical significance was defined when p < 0.05 (*p < 0.05, **p < 0.01, ***p < 0.001, ****p < 0.0001).

## List of Supplementary Materials

Fig S1 to S7

Tables S1 to S3

## Acknowledgments

We acknowledge the facilities at the Harry Perkins Institute of Medical Research, the scientific and technical assistance of Kevin Li with flow cytometry. We thank Harry Perkins Institute of Medical Research Bioresources staff for their assistance with animal husbandry. Part of this research was undertaken on the XFM beamline at the Australian Synchrotron, part of ANSTO. This study makes use of data generated by the MOSAIC consortium (Owkin, Charité – Universitätsmedizin Berlin, Lausanne University Hospital - CHUV, Erlangen Hospital, Gustave Roussy Institute, University of Pittsburgh) and made available through the MOSAIC Window initiative. MOSAIC consortium bears no responsibility for the further analysis or interpretation of these data beyond what published by the MOSAIC consortium partners. Silicon nitride windows were manufactured by the Melbourne Centre for Nanofabrication (MCN) in the Victorian node of the Australian National Fabrication Facility (ANFF).

## Funding

WA Department of Health Future Health Research and Innovation Fund Translation Fellowship (JC)

UWA Australia Research Collaboration Award Cancer Council WA (JC)

iCare Dust Diseases Board Project Grant

NHMRC Centres of Research Excellence APP1197652 (National Centre for Asbestos Related Diseases)

Australian Government Research Training Programme scholarship supported by an AINSE Ltd Postgraduate Research Award (KRIS)

Australian Synchrotron Merit Access programme AS252/XFM/23805; AS243/XFM/22244

SPHERE Cancer Clinical Academic Group Fellowship supported by Cancer Institute NSW Capacity Building Grant (BK)

## Author contributions

Conceptualization: OV, JC

Methodology OV, JC, AC, RMG, DN, AC, DH, LB, WJL, WLC

Investigation: KLPS, ND, CP, ALP, NP, GC, JR, SW, WSY, KRIS

Visualization: KLPS, NP, RR, ND, VP, JC

Funding acquisition: OV, JC, WLC, RMG

Project administration: JC, OV, KLPS

Supervision: OV, DN, JC, KLPS

Writing – original draft: JC, ND, KLPS

Writing – review & editing: all authors

## Data and materials availability

All data associated with this study are present in the paper or the Supplementary Materials. Sequencing data will be deposited on GEO upon paper acceptance.

