## Supplementary Figures 1-7; Table 1 for "Copper chelation reprograms the tumour microenvironment to enhance immune checkpoint blockade therapy in preclinical mesothelioma"

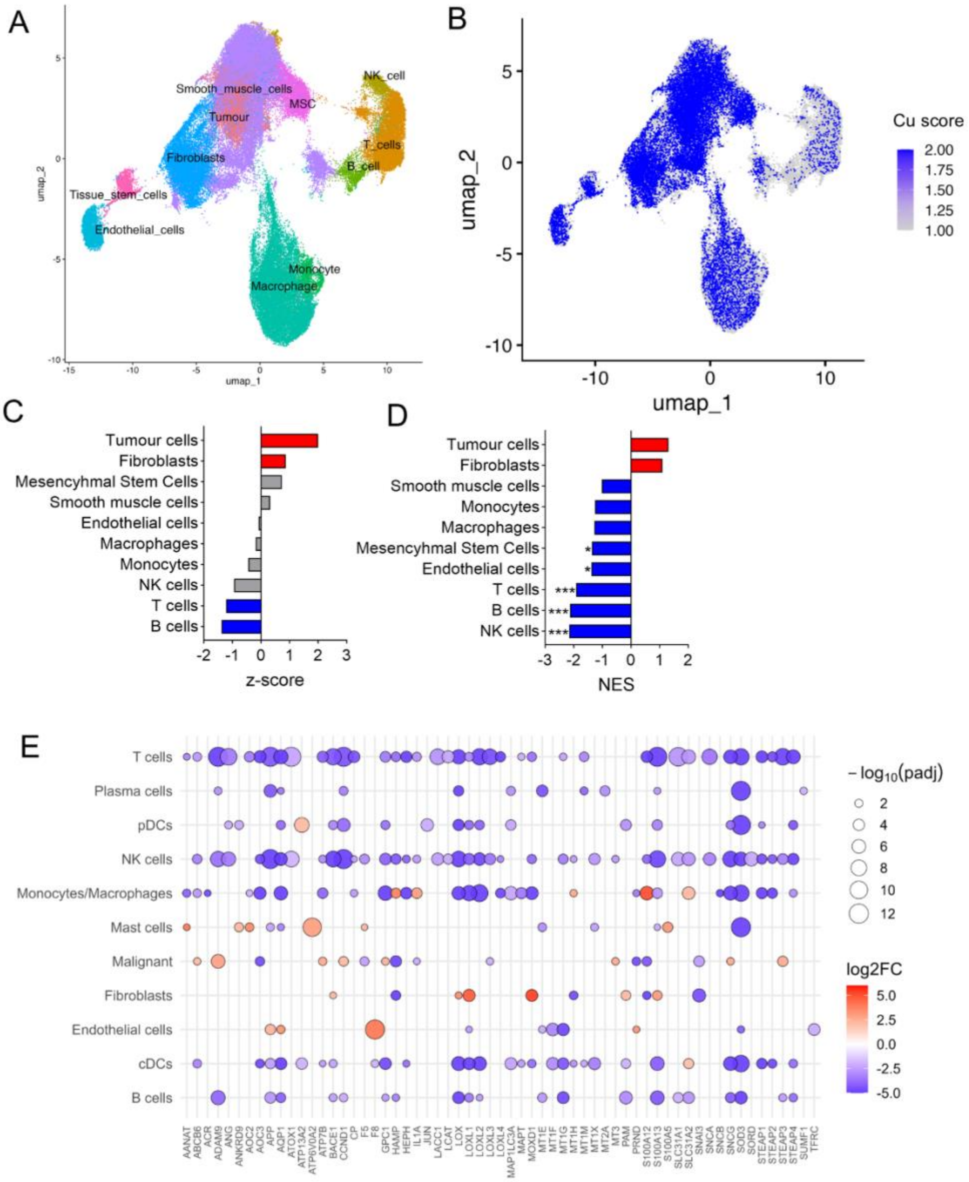


**Supplementary Fig. 1. Single cell RNAseq analysis of a cohort of human mesothelioma samples.** (**A**) UMAP projection of single-cell RNA-seq data from human mesothelioma samples (MOSAIC-MESO n = 10), annotated with major cell populations (top) and overlaid with (**B**) per-cell copper homeostasis gene-set score. (**C**) Average expression of copper homeostasis gene set for indicated cell subsets in MOSAIC-MSO, shown as z-scores. (**D**) Normalised enrichment scores (NES) from fgsea of copper homeostasis gene sets across cell subsets. (**E**) Plot displaying differential expressed genes (DEGs) of copper homeostasis genes across major cell types in Giotti et al. DEGs with p-adj < 0.05 and log2FC > 1 displayed.

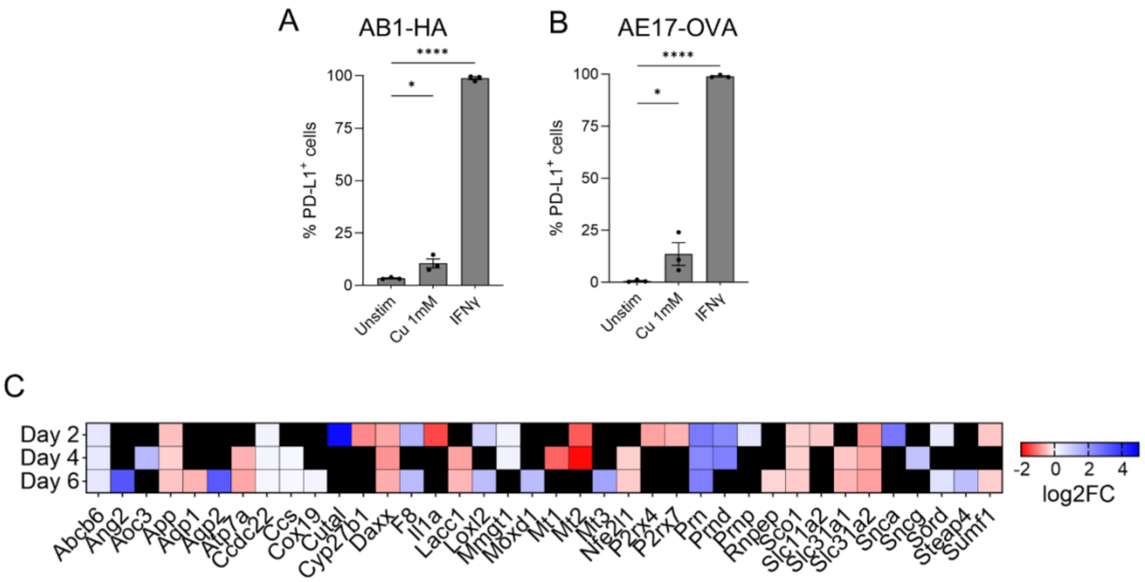


**Supplementary Fig. 2. Copper treatment upregulates PD-L1 on mesothelioma tumour cell lines.** Dot plot displaying percentage of (**A**) AB1-HA and (**B**) AE17-OVA tumour cells expressing PD-L1 after 4 hours of treatment with 1 mM $CuCl_2$, or 200 U IFNγ *in vitro*, measured by flow cytometry. One-way ANOVA with Benjamini–Hochberg correction for multiple comparisons used to compare treatment groups. (n = 3/group; *$p < 0.05$, ****$p < 0.001$). (**C**) Heatmap of copper homeostasis genes with significant response × timepoint interaction effects during ICB treatment (padj < 0.05). Colours represent interaction log2 fold-change values at each post-ICB timepoint, with red indicating greater relative temporal change in non-responders and blue indicating greater relative temporal change in responders. Black denotes non-significant gene timepoint comparisons.

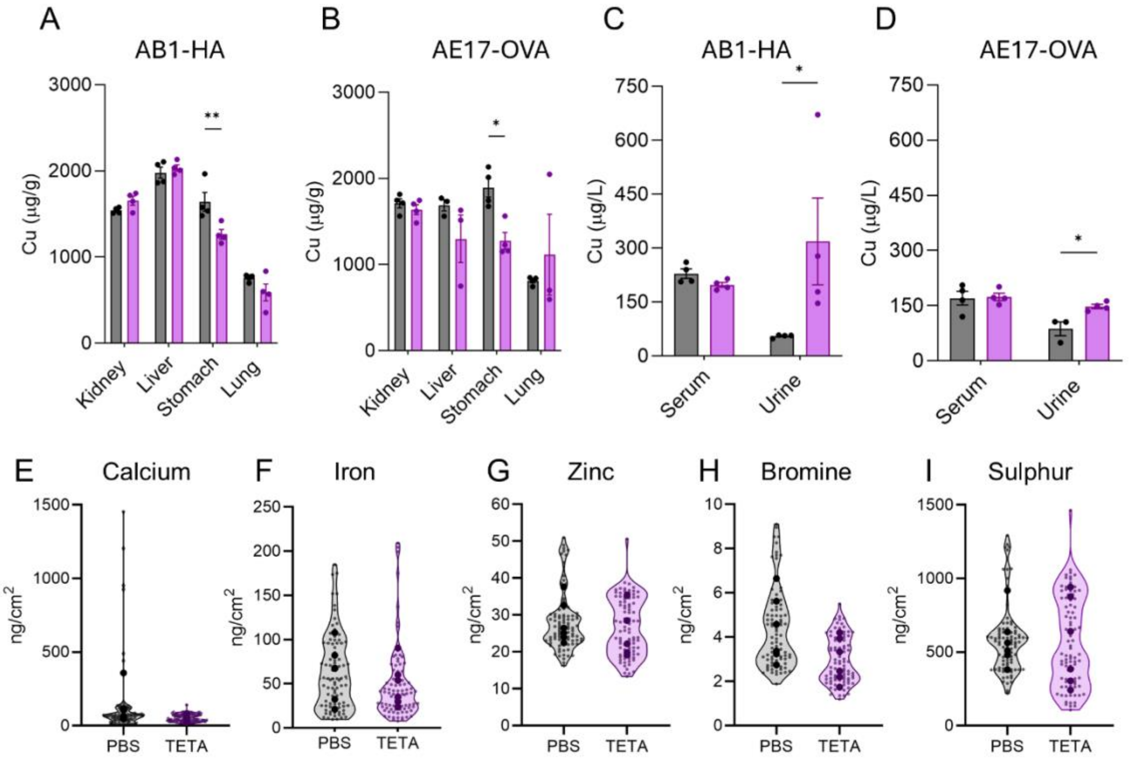


**Supplementary Fig. 3. Effects of TETA on systemic copper and intratumoural trace elements.** Atomic Absorption Spectrophotometry copper measurements in (**A**, **B**) organs (kidney, liver, stomach and lung), (**C**, **D**) serum and urine from AB1-HA or AE17-OVA mice treated with 7 days vehicle (PBS) or TETA. Data presented as mean ± SEM (n = 4/group) and two independent experiments. Two-tailed paired t-test used to compare differences between TETA and PBS for individual organs. XFM measurements of intratumoural (**E**) calcium, (**F**) iron, (**G**) zinc, (**H**) bromine and (**I**) sulphur in AB1-HA and AE17-OVA tumours from PBS and TETA treated animals. Violin plots display the distribution from individual measurements from multiple regions per treatment (n = 6/group), nested within individual biological samples. Nested t-test used to compare differences between treatment groups for XFM data. *p < 0.05, **p < 0.01.

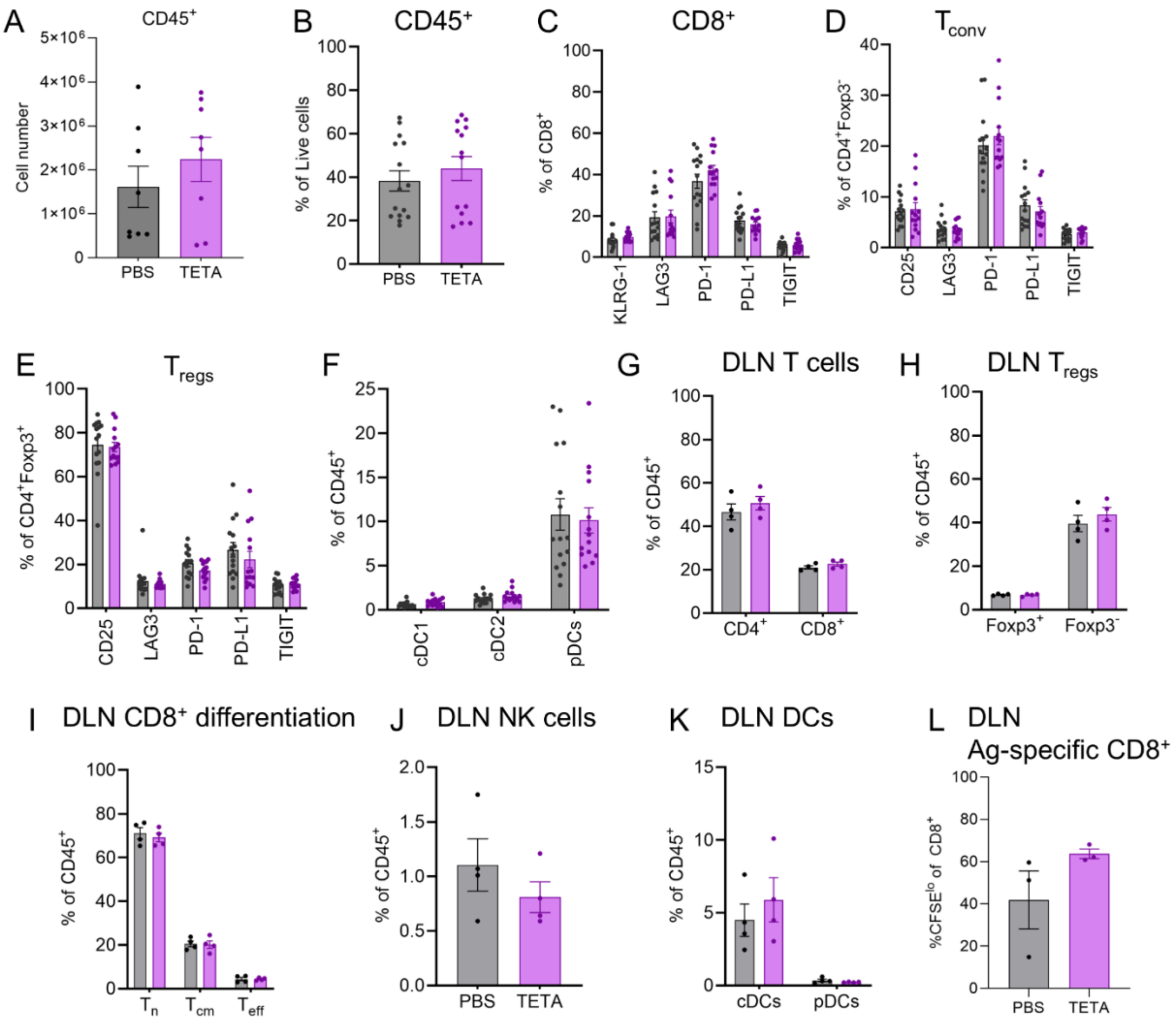


**Supplementary Fig. 4. Tumour infiltrating and draining lymph node cells with TETA treatment.** Dot plots displaying (**A**) number and (**B**) proportion of CD45$^+$ cells, percentage of (**C**) CD8$^+$, (**D**) CD4$^+$Foxp3$^-$ (T$_{conv}$), (**E**) CD4$^+$Foxp3$^+$ (T$_{regs}$) expressing activation and inhibitory receptors, and (**F**) percentage of cDC1, cDC2 and pDCs in AB1-HA s.c tumours, TETA versus PBS (n = 8-15/ group). Dot plots displaying percentages of (**G**) CD4$^+$, CD8$^+$, (**H**) CD4$^+$Foxp3$^{+/-}$, (**I**) naïve (CD44$^{lo}$CD62L$^{hi}$), central memory (CD44$^{hi}$CD62L$^{hi}$) and effector (CD44$^{hi}$CD62$^{lo}$) CD8$^+$ T cells, (**J**) CD335$^+$ NK cells, (**K**) DCs, and (**L**) proliferating (CFSE$^{lo}$), transferred HA-specific TCR transgenic T cells in tumour draining lymph nodes (DLN), TETA versus PBS (n = 3-4/group).

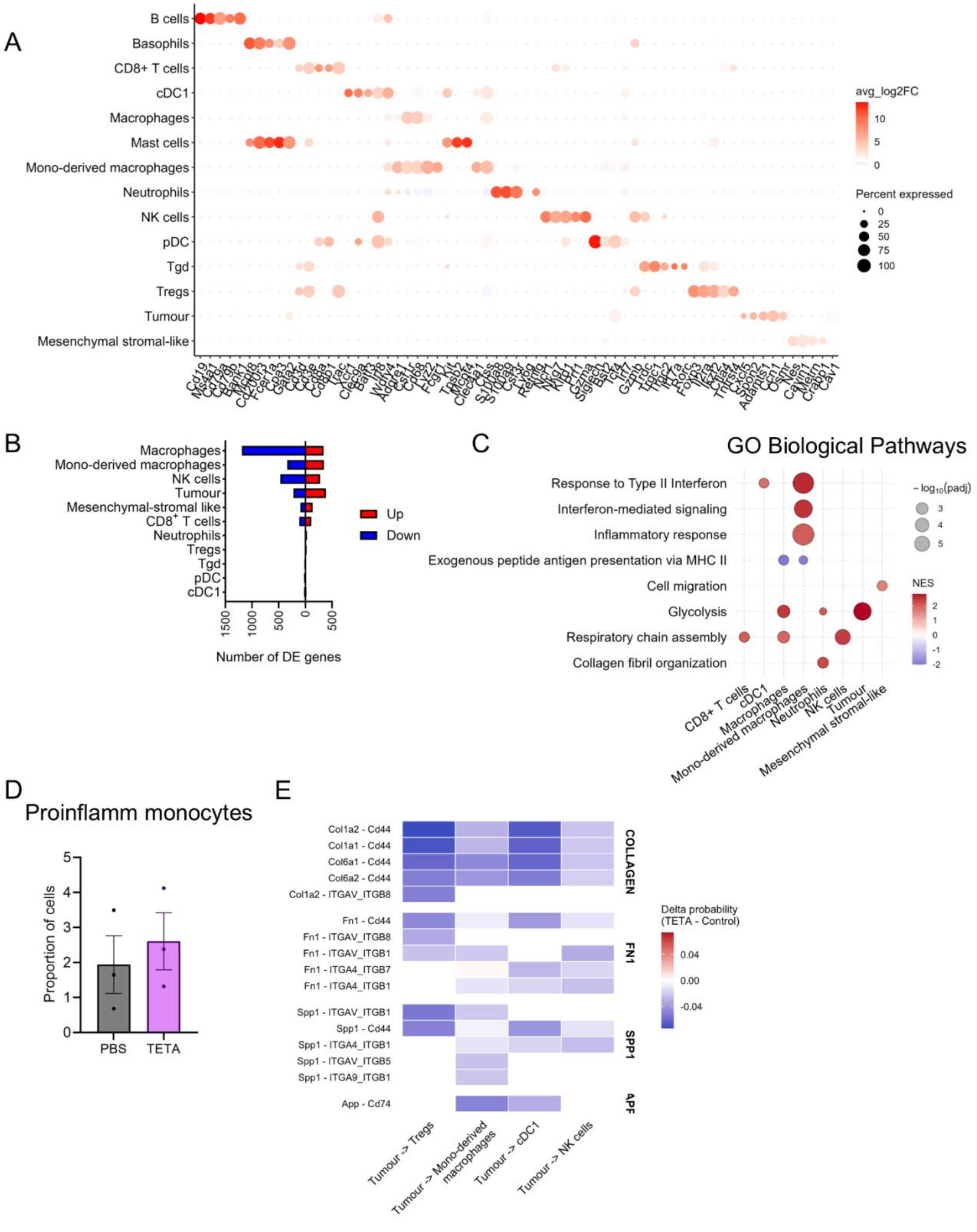


**Supplementary Fig. 5. TETA treatment reprograming of the tumour transcriptome.** (**A**) Dot plot of gene expression markers used to classify the cell clusters in main Figure 4. (**B**)

Number of differentially expressed genes (DEGs) for each cell cluster in TETA treated animals relative to PBS. (**C**) Plot depicting GSEA for GO Biological Processes pathways using the fgsea package. Significantly enriched pathways in TETA-treated (red) or PBS-treated (blue) (padj < 0.05) across individual cell clusters. (**D**) Dot plot representing proportion of cells that express proinflammatory monocyte signature. (**E**) Heatmap of inferred ligand–receptor interaction probabilities from tumour compartments to immune populations, demonstrating decreased extracellular matrix–related signalling following TETA treatment.

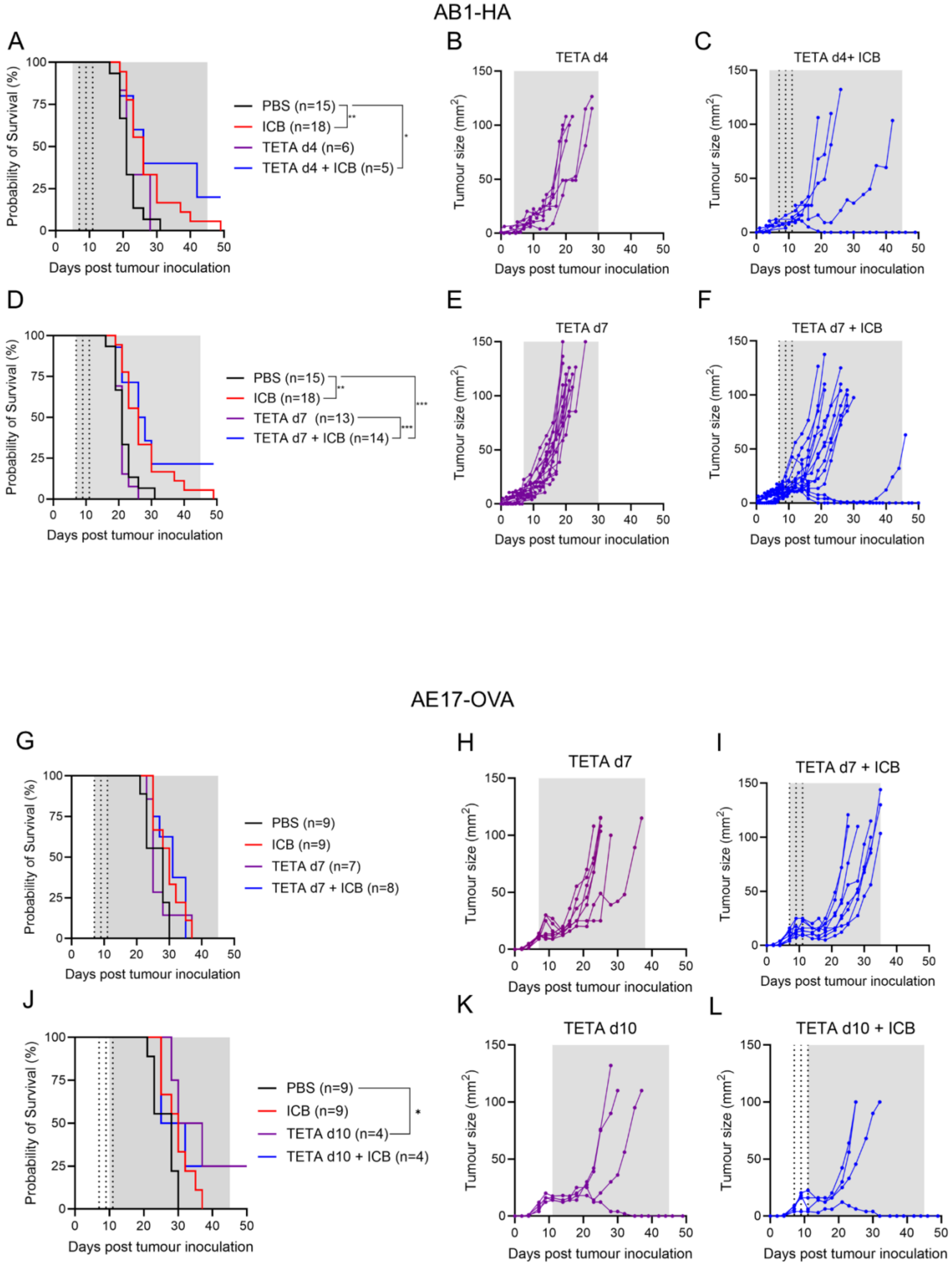
AB1-HA
A
Probability of Survival (%)
Days post tumour inoculation
PBS (n=15)
ICB (n=18)
TETA d4 (n=6)
TETA d4 + ICB (n=5)
B
TETA d4
C
TETA d4+ ICB
Tumour size (mm$^2$)
D
PBS (n=15)
ICB (n=18)
TETA d7 (n=13)
TETA d7 + ICB (n=14)
E
TETA d7
F
TETA d7 + ICB
AE17-OVA
G
PBS (n=9)
ICB (n=9)
TETA d7 (n=7)
TETA d7 + ICB (n=8)
H
TETA d7
I
TETA d7 + ICB
J
PBS (n=9)
ICB (n=9)
TETA d10 (n=4)
TETA d10 + ICB (n=4)
K
TETA d10
L
TETA d10 + ICB

**Supplementary Fig. 6. Schedule-dependent effects of TETA and ICB on tumour growth and survival in mesothelioma-bearing mice.** (**A**) Kaplan-Meier survival curves of subcutaneous AB1-HA tumours in mice treated with vehicle (PBS, black), immune checkpoint blockade (ICB; anti-CTLA-4 and anti- PD-1; red), TETA (purple) and combination TETA and ICB (blue), with TETA initiated at day 4 (d4) post tumour inoculation. Individual tumour growth curves for (**B**) TETA (d4), (**C**) TETA (d4) + ICB. (**D**) Kaplan–Meier survival curves of subcutaneous AB1-HA tumours treated with PBS, ICB, TETA (d7) and combination TETA (d7) and ICB. Individual tumour growth curves for (**E**) TETA (d7), (**F**) TETA (d7) + ICB. (**G**) Kaplan–Meier survival curves of subcutaneous AE17-OVA tumours treated with PBS, ICB, TETA, and combination TETA and ICB, with TETA initiated at day 7 (d7) after tumour inoculation. Individual tumour growth curves for (**H**) TETA (d7) and (**I**) TETA (d7) + ICB. (**J**) Kaplan–Meier survival curves of subcutaneous AE17-OVA tumours treated with PBS, ICB, TETA, and combination TETA and ICB.  TETA was initiated at d4 after tumour inoculation. Individual tumour growth curves for (**K**) TETA (d7) and (**L**) TETA (d7) + ICB. Pairwise comparisons for survival were performed using a Mantel–Cox log-rank test. Dotted vertical lines denote ICB treatment time points with grey shaded areas indicating TETA treatment. Group sizes indicated in figures.

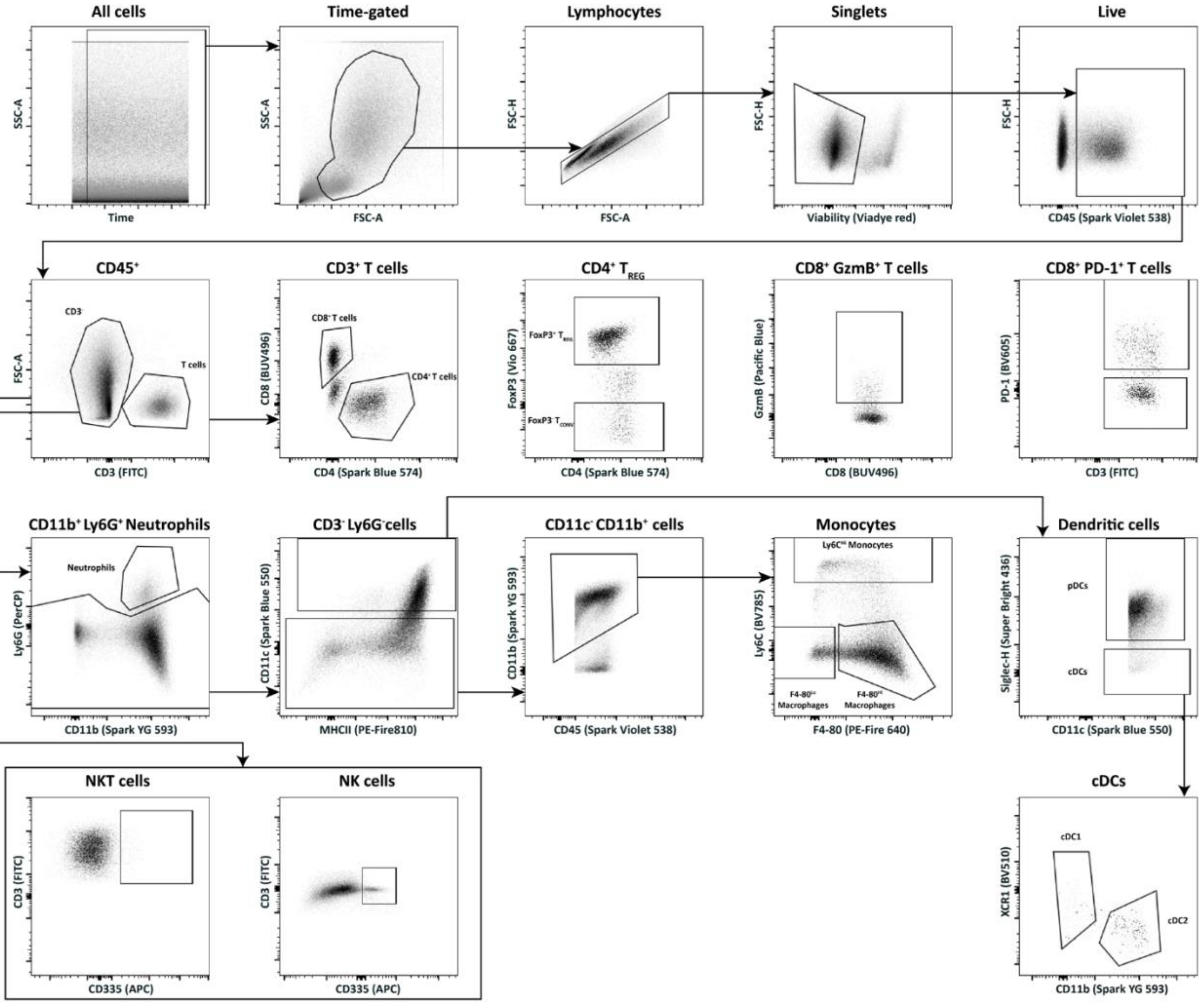


**Supplementary Fig. 7. Flow cytometry gating strategy for tumour infiltrating lymphoid and myeloid cells.**

**Supplementary Table 1. Significantly enriched copper genes (tumour versus normal tissue).**

| Gene | Median Diff | p_adj |
|---|---|---|
| KBP4 | 0.72 | 1.39E-16 |
| LOX | 0.65 | 2.77E-16 |
| SP1 | 0.63 | 1.39E-16 |
| BECN1 | 0.58 | 1.39E-16 |
| GPC1 | 0.57 | 1.39E-16 |
| ATP6V0A2 | 0.57 | 1.39E-16 |
| STEAP3 | 0.57 | 1.39E-16 |
| MDM2 | 0.55 | 1.39E-16 |
| XIAP | 0.51 | 1.39E-16 |
| LOXL1 | 0.51 | 5.18E-16 |
| ARF1 | 0.47 | 1.39E-16 |
| NFE2L2 | 0.46 | 1.39E-16 |
| HEPH | 0.45 | 1.39E-16 |
| TP53 | 0.43 | 1.39E-16 |
| COX17 | 0.42 | 1.39E-16 |
| AP1S1 | 0.41 | 1.39E-16 |
| LOXL2 | 0.41 | 1.39E-16 |
| ADAM9 | 0.41 | 7.51E-16 |
| SNCG | 0.37 | 1.39E-16 |
| ATP13A2 | 0.36 | 1.39E-16 |
| STEAP4 | 0.34 | 2.61E-13 |
| AQP1 | 0.33 | 1.08E-14 |
| DAXX | 0.33 | 1.39E-16 |
| AP1B1 | 0.33 | 1.39E-16 |
| SCO1 | 0.32 | 1.90E-16 |
| HSF1 | 0.31 | 1.39E-16 |
| CCDC22 | 0.29 | 1.57E-16 |
| CDK1 | 0.29 | 1.31E-11 |
| HSPA5 | 0.27 | 5.61E-15 |
| GSK3B | 0.25 | 8.50E-16 |
| CCND1 | 0.24 | 4.52E-11 |
| ANKRD9 | 0.24 | 5.86E-16 |
| SIL1 | 0.22 | 1.15E-14 |
| AKT1 | 0.21 | 2.56E-15 |
| SLC31A1 | 0.21 | 2.20E-13 |
| PTEN | 0.21 | 1.15E-14 |
| ATP7A | 0.21 | 4.67E-10 |
| ADNP | 0.21 | 4.30E-12 |
| ATF6 | 0.21 | 1.39E-16 |
| MOXD1 | 0.19 | 5.95E-10 |
| CASP3 | 0.19 | 7.13E-11 |
| BACE1 | 0.19 | 1.64E-14 |
| CCS | 0.18 | 2.60E-16 |
| COX11 | 0.18 | 6.72E-15 |
| SEC61G | 0.17 | 2.80E-12 |
| ATOX1 | 0.17 | 9.61E-15 |
| APP | 0.17 | 9.93E-13 |
| LCAT | 0.17 | 5.95E-10 |
| LACC1 | 0.16 | 3.85E-10 |
| ATP6AP1 | 0.16 | 3.47E-12 |
| STEAP1 | 0.15 | 1.87E-03 |
| ADAM10 | 0.15 | 5.15E-10 |
| ADAM17 | 0.15 | 2.50E-07 |
| HAMP | 0.14 | 5.94E-04 |
| RNF7 | 0.14 | 8.72E-11 |
| STEAP2 | 0.13 | 3.96E-02 |
| APC | 0.13 | 3.18E-07 |
| GSS | 0.13 | 3.31E-14 |
| CUTA | 0.12 | 2.36E-10 |
| COX19 | 0.12 | 9.04E-16 |
| CDKN2A | 0.11 | 7.81E-05 |
| ATP7B | 0.10 | 6.27E-06 |
| COMMD1 | 0.09 | 9.26E-09 |
| PIK3CA | 0.08 | 1.00E-06 |
| MT2A | 0.08 | 5.00E-04 |
| P2RX4 | 0.07 | 5.14E-07 |
| SUMF1 | 0.07 | 5.29E-05 |
| CUTC | 0.07 | 5.00E-04 |
| MTF2 | 0.07 | 4.50E-03 |
| COG2 | 0.07 | 4.03E-03 |
| SLC11A2 | 0.06 | 1.11E-05 |
| LOXL4 | 0.05 | 1.51E-02 |
| XBP1 | 0.04 | 1.91E-02 |
| JUN | 0.04 | 1.32E-02 |
| MTF1 | 0.03 | 4.55E-03 |
| SOD1 | 0.03 | 5.14E-03 |
| SOD3 | -0.07 | 3.44E-02 |
| FOXO3 | -0.07 | 1.28E-05 |
| MT1F | -0.08 | 2.91E-03 |
| F8 | -0.08 | 5.39E-03 |
| S100A13 | -0.10 | 5.23E-06 |
| XAF1 | -0.10 | 1.19E-02 |
| MT1G | -0.12 | 3.89E-07 |
| SNCA | -0.18 | 1.40E-05 |
| AOC2 | -0.18 | 1.47E-09 |
| PRND | -0.20 | 4.51E-04 |
| ANG | -0.23 | 1.94E-10 |
| MT1M | -0.25 | 2.47E-04 |
| MT1X | -0.26 | 3.50E-11 |
| AOC1 | -0.28 | 1.58E-10 |

| | | | | | |
|---|---|---|---|---|---|
| AANAT | -0.30 | 1.98E-11 | SLC31A2 | -0.56 | 1.79E-16 |
| MT3 | -0.32 | 1.08E-09 | SNCB | -0.63 | 2.07E-14 |
| AOC3 | -0.33 | 4.31E-08 | ACR | -0.63 | 1.46E-14 |
| S100A5 | -0.33 | 2.58E-09 | S100A12 | -0.67 | 2.78E-14 |
| ALB | -0.34 | 9.50E-11 | TYR | -0.72 | 4.15E-14 |
| IL1A | -0.38 | 1.17E-12 | CA6 | -0.86 | 1.39E-16 |
| MT1H | -0.39 | 1.81E-15 | AQP2 | -0.97 | 1.39E-16 |
| DCT | -0.39 | 1.66E-10 | CYP1A1 | -1.00 | 1.39E-16 |
| CP | -0.47 | 7.89E-11 | MT4 | -1.10 | 1.39E-16 |
| DBH | -0.48 | 2.09E-11 | MT1HL1 | -1.14 | 1.39E-16 |
| TMPRSS6 | -0.50 | 3.79E-16 | APOA4 | -1.15 | 1.39E-16 |
| SCO2 | -0.53 | 1.48E-16 | | | |

**Supplementary Table 2. Copper homeostasis genes associated with better median overall survival in TCGA-MESO.**

| Gene | Padj |
|---|---|
| ACR | 3.00E-02 |
| ADAM10 | 7.52E-03 |
| ADAM17 | 1.57E-02 |
| ADAM9 | 2.29E-02 |
| AQP2 | 6.46E-03 |
| ATP6V0A2 | 3.05E-02 |
| CCND1 | 2.68E-03 |
| CDK1 | 3.42E-06 |
| GPC1 | 3.70E-02 |
| LCAT | 3.29E-02 |
| LOX | 4.85E-03 |
| LOXL1 | 4.10E-03 |
| LOXL2 | 2.89E-06 |
| PRND | 4.60E-02 |
| S100A13 | 1.40E-02 |
| SLC31A1 | 1.53E-03 |
| SORD | 1.10E-02 |
| STEAP3 | 2.27E-02 |
| ANKRD9 | 3.69E-02 |
| APOA4 | 3.22E-02 |
| ATP7A | 2.89E-02 |
| HAMP | 3.79E-02 |
| MT4 | 1.34E-02 |
| NFE2L2 | 4.88E-02 |
| STEAP2 | 2.53E-02 |

Padj, adjusted p-value from log-rank tests comparing high vs low expression groups (median split). Orange shading indicates genes for which high tumour expression is associated with improved survival, and blue shading indicates genes for which low tumour expression is associated with improved survival.

**Supplementary Table 3. Panel of antibodies used for spectral flow cytometry.**

| Target | Fluor | Antigen | Dilution | Clone | Company | Cat# |
|---|---|---|---|---|---|---|
| Dead cells | ViaDye ™ Red | NA | 1:20000 | NA | Cytek Biosciences | SKU R7-60008 |
| CD45 | Spark Violet 538 | CD45 | 1:250 | 30-F11 | Biolegend | 103179 |
| CD3 | FITC | CD3 | 1:500 | 17A2 | Cytek Biosciences | 35-0032-U100 |
| CD8 | BUV496 | CD8 | 1:250 | 53-6.7 | BD | 750024 |
| CD4 | Spark Blue 574 | CD4 | 1:1000 | GK1.5 | Biolegend | 100489 |
| FOXP3 | Vio R667 | Foxp3 | 1:100 | REA788 | Miltenyi Biotech | 130-111-682 |
| CD25 | BV480 | CD25 | 1:500 | PC61.5 | BD | 566202 |
| Granzyme B | Pacific Blue | GzmB | 1:250 | GB11 | Biolegend | 515407 |
| CD44 | Alexa Fluor 700 | CD44 | 1:1000 | IM7 | Thermo | 56-0441-80 |
| CD62L (L-Selectin) | BV570 | CD62L | 1:500 | MEL-14 | Biolegend | 104433 |
| KLRG1 | APC-eFluor 780 | KLRG1 | 1:1000 | 2F1 | Thermo | 47-5893-80 |
| CD223 | BUV737 | LAG-3 (CD223) | 1:500 | C9B7W | BD | 741820 |
| PD-L1 | PerCP-eFluor710 | PD-L1 (CD274) | 1:500 | MIH5 | Thermo | 46-5982-82 |
| CD279 | BV605 | PD-1 (CD279) | 1:500 | 29F.1A12 | Biolegend | 135219 |
| TIGIT | PE-Dazzle 594 | TIGIT | 1:250 | 1G9 | Biolegend | 142109 |
| CD335 (NKp46) | APC | CD335 (NKp46) | 1:250 | 29A1.4 | Biolegend | 137607 |
| Ly-6G | PerCP | Ly6G | 1:1000 | 1A8 | Biolegend | 127653 |
| Ly-6C | BV785 | Ly6C | 1:1000 | HK1.4 | Biolegend | 128041 |

| | | | | | | |
|---|---|---|---|---|---|---|
| Siglec-H | Super Bright 436 | SiglecH | 1:250 | 440c | Thermo | 62-0333-82 |
| XCR1 | BV510 | XCR1 | 1:100 | ZET | Biolegend | 148218 |
| F4-80 | PE-Fire 640 | F4-80 | 1:250 | QA17A29 | Biolegend | 157319 |
| CD11b | Spark YG 593 | CD11b | 1:400 | M1/70 | Biolegend | 101281 |
| CD11c | Spark Blue 550 | CD11c | 1:1000 | N418 | Biolegend | 117365 |
| MHC II (I-A+I-E) | PE-Fire 810 | MHC-II | 1:400 | M5/114.15.2 | Biolegend | 107667 |
